# *Abcg2*-Expressing Side Population Cells Contribute to Cardiomyocyte Renewal through Fusion

**DOI:** 10.1101/690339

**Authors:** Amritha Yellamilli, Yi Ren, Ron T. McElmurry, Jonathan P. Lambert, Polina Gross, Sadia Mohsin, Steven R. Houser, John W. Elrod, Jakub Tolar, Daniel J. Garry, Jop H. van Berlo

## Abstract

The adult mammalian heart has a limited regenerative capacity. Therefore, identification of <u>endogenous</u> cells and mechanisms that contribute to cardiac regeneration is essential for the development of targeted regenerative therapies. The side population (SP) phenotype enriches for stem cells throughout the body, and SP cells have been identified in the heart. We generated a novel Abcg2-driven, genetic lineage-tracing mouse model and show efficient labeling of SP cells. Labeled SP cells give rise to terminally differentiated cells in bone marrow and intestines. In the heart, we find that Abcg2-expressing SP cells contribute to cardiomyocyte labeling, which is further enhanced in response to different forms of cardiac injury. We find that cardiac SP cells fuse with preexisting cardiomyocytes, which stimulates cardiomyocyte cell cycle entry and likely proliferation, instead of directly differentiating into cardiomyocytes. Our findings provide evidence that cardiac SP cells contribute to endogenous cardiac regeneration through cardiomyocyte fusion.

## INTRODUCTION

A critical feature of heart failure that limits the effectiveness of current therapies is loss of functional cardiomyocytes(Chiong et al., 2011; van Empel et al., 2005). Since the adult heart has a limited ability to generate new cardiomyocytes, strategies that enhance cardiomyocyte proliferation in failing hearts are actively being developed(Tzahor & Poss, 2017; van Berlo & Molkentin, 2014). The discovery of progenitor cells in adult mammalian hearts initially sparked excitement about targeting these cells to regenerate the heart; however, recent studies have caused the field of cardiac regeneration to shift its focus away from progenitor cells as a direct source of new cardiomyocytes(Beltrami et al., 2003; Hong et al., 2014; Liu et al., 2016; Sanganalmath & Bolli, 2013; Sultana et al., 2015; van Berlo et al., 2014). Currently, proliferation of preexisting cardiomyocytes is believed to be the sole source of new cardiomyocytes, however, it is unclear if other sources of new cardiomyocytes exist that can be targeted for new regenerative therapies(Eschenhagen et al., 2017; Senyo et al., 2013).

To assess whether endogenous cardiac progenitor cells exist in an unbiased manner without using previously studied membrane markers, we utilized the side population phenotype that enriches for stem and progenitor cells throughout the body and in different forms of cancer(Golebiewska, Brons, Bjerkvig, & Niclou, 2011; Hierlihy, Seale, Lobe, Rudnicki, & Megeney, 2002; Martin et al., 2004; Unno, Jain, & Liao, 2012). The side population phenotype is the ability of stem and progenitor cells to extrude Hoechst 33342, a fluorescent DNA dye, out of their cytoplasm. To identify side population cells (SPCs), a single cell suspension is incubated with Hoechst 33342 and analyzed with flow cytometry to identify cells that extrude Hoechst 33342 and thereby sort to the “side” of the main population of cells(Goodell, Brose, Paradis, Conner, & Mulligan, 1996). SPCs were first described as a way to enrich for hematopoietic stem cells capable of long-term bone marrow reconstitution after transplantation into lethally irradiated mice(Goodell et al., 1996). Since then, SPCs have been identified in intestines, skeletal muscle, and the heart(Yellamilli & van Berlo, 2016).

Isolated cardiac side population cells (cSPCs) are enriched for cells that self-renew and differentiate into multiple cardiac lineages in cell culture and after transplantation. However, it is unknown whether the side population phenotype enriches for endogenous cardiac progenitor cells that can give rise to cardiomyocytes(Noseda et al., 2015; Oyama et al., 2007; Unno et al., 2012; Yellamilli & van Berlo, 2016). Cultured cSPCs form colonies at ten times the rate of other non-cardiomyocytes isolated from the heart(Martin et al., 2004; Pfister et al., 2005). Primary and secondary cSPC clones can be maintained in cell culture for over ten months with preservation of the side population phenotype and without undergoing replicative senescence(Noseda et al., 2015). In cell culture and transplantation studies, cSPCs are also multipotent. They differentiate towards cardiomyocyte, endothelial cell or smooth muscle cell lineages under specific culture conditions(Yellamilli & van Berlo, 2016). In preclinical studies, transplanted cSPCs engraft in the heart after cryoinjury or myocardial ischemia (MI) and give rise to cardiomyocytes, endothelial cells, smooth muscle cells and fibroblasts(Liang, Tan, Gaudry, & Chong, 2010; Noseda et al., 2015; Oyama et al., 2007). Notably, cSPCs do not express *Kit* and have a distinct expression pattern from cardiac *Kit*-expressing cells(Dey et al., 2013; Noseda et al., 2015; Pfister et al., 2005). Studies of isolated cSPCs are limited to the context of cell culture and transplantation so whether the side population phenotype can be used to enrich for endogenous cardiac progenitor cells that are crucial for maintaining cardiac homeostasis and regeneration is unclear. Recent publications have also questioned the existence of any type of progenitor cell that can differentiate into cardiomyocytes in the adult heart(He et al., 2017; Kretzschmar et al., 2018; Li et al., 2018). Here, we assessed whether cSPCs are enriched for endogenous cardiac progenitor cells, and generated a lineage-tracing mouse model driven by the *Abcg2* gene, which encodes a transporter essential for the side population phenotype(Krishnamurthy & Schuetz, 2006; Martin et al., 2004; Pfister et al., 2008; Zhou et al., 2002; Zhou et al., 2001).

## RESULTS

### Lineage-tracing mouse model reliably and specifically labels Abcg2-expressing cells

To lineage-trace endogenous SPCs, we generated an *Abcg2*-driven lineage-tracing mouse model in which cDNA encoding a tamoxifen-inducible MerCreMer (MCM) fusion protein was inserted into the endogenous *Abcg2* locus (Abcg2^MCM/+^). These mice were cross-bred with Cre-inducible reporter mice, R26^GFP/GFP^, to generate experimental Abcg2^MCM/+^ R26^GFP/+^ mice (Figure 1A). To induce recombination, two different tamoxifen injection strategies were used (Figures 1B and 1C). First, we verified that *Abcg2*-expressing cells were specifically labeled in Abcg2^MCM/+^ R26^GFP/+^ mice injected with tamoxifen. ABCG2 is expressed at the apical surfaces of renal proximal tubule and intestinal epithelium, on canalicular membranes of hepatocytes and by cardiac endothelium(Dekaney, Rodriguez, Graul, & Henning, 2005; Higashikuni et al., 2010; Horsey, Cox, Sarwat, & Kerr, 2016). We confirmed this reported expression pattern with immunofluorescent staining using an ABCG2-specific antibody on renal, ileal, hepatic and cardiac tissue sections from wild type mice (Figure 1D). No staining was observed on sections from *Abcg2*-knockout mice(Zhou et al., 2002), confirming specificity of the antibody used (Figure S1A). To determine whether the *Abcg2*-driven lineage-tracing mouse model labels *Abcg2*-expressing cells, we imaged native fluorescence of GFP in tissues harvested from tamoxifen-injected Abgc2^MCM/+^ R26^GFP/+^ mice at week 9 (Figure 1B). We observed extensive labeling of cells in the kidney, ileum, liver and heart known to express ABCG2 (Figure 1E). Notably, no GFP-labeling was observed in vehicle-injected Abgc2^MCM/+^ R26^GFP/+^ mice, indicating no leaky Cre activity (Figure S1B).GFP-labeling in other organs appeared to be restricted to endothelial cells (Figure S1C).

**Figure 1.**
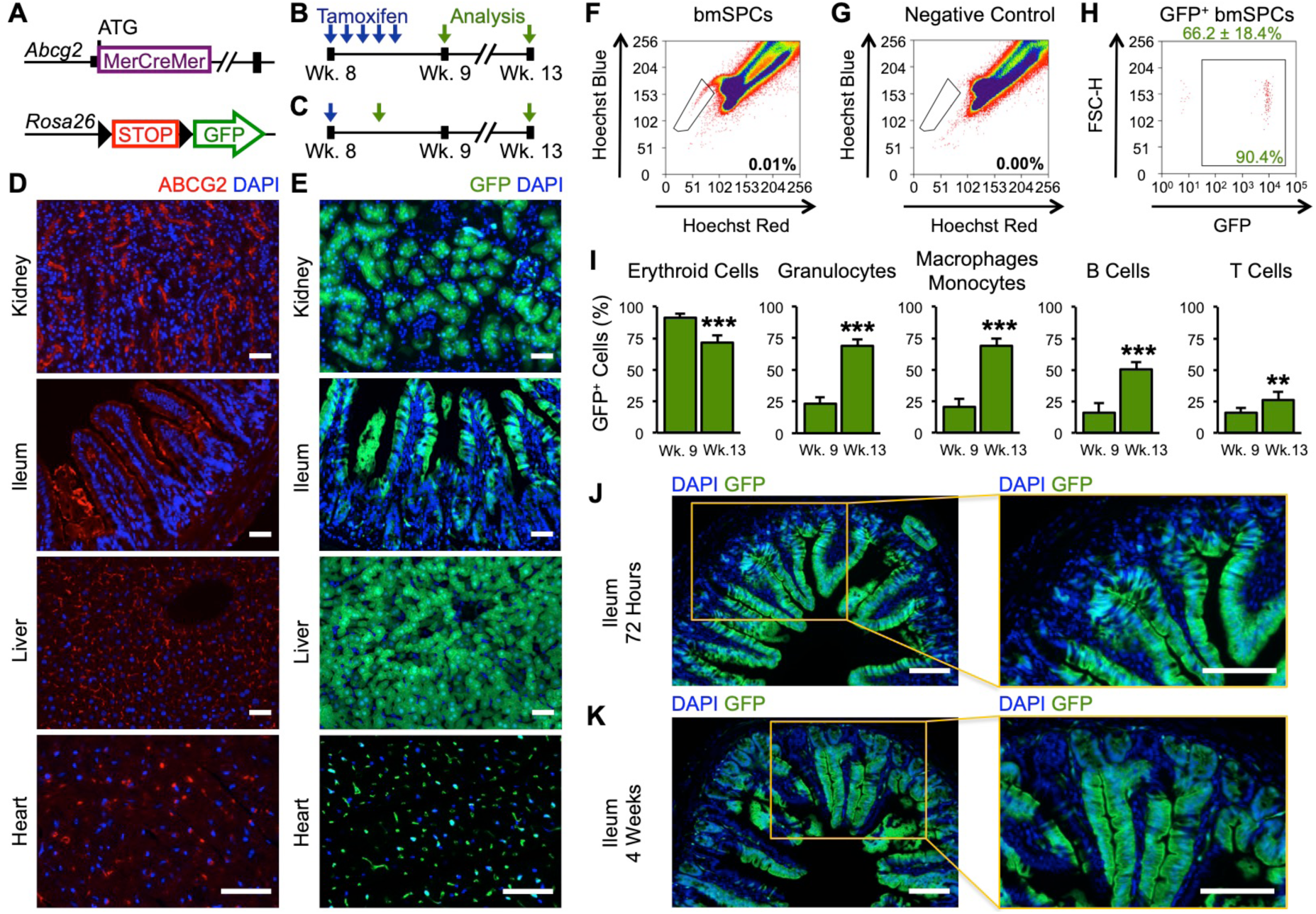
Bone marrow SPCs and intestinal stem cells give rise to differentiated lineages in vivo. **(A)** Genetics of experimental Abcg2^MCM/+^ R26^GFP/+^ mice. **(B)** Experimental timeline used to induce recombination in bone marrow and cardiac studies. Blue arrows represent an intraperitoneal injection of tamoxifen. Green arrows represent time points at which GFP-labeling was assessed 72 hours (week 9) and 4 weeks (week 13) after the fifth tamoxifen injection. **(C)** Experimental timeline used to induce recombination in intestinal studies. GFP-labeling was assessed 72-hours and 4-weeks after single tamoxifen injections. **(D)** Fluorescent images of sections from C57Bl/6J mice stained for ABCG2. **(E)** Images of native GFP fluorescence of sections from tamoxifen-injected Abcg2^MCM/+^ R26^GFP/+^ mice at week 9. **(F)** Flow cytometry plot of bone marrow SPCs (bmSPCs) in tamoxifen-injected Abcg2^MCM/+^ R26^GFP/+^ mice (box to the left of the main population, number inside FACS plot is the percent of bmSPCs for this sample, n=6). **(G)** Corresponding flow cytometry plot of bmSPC negative control stained Verapamil, which ensures accurate identification of bmSPCs (number inside FACS plot is percent of SPCs for this sample, n=6). **(H)** Flow cytometry plot of GFP fluorescence of bmSPCs (number within gate is the percent of GFP^+^ bmSPCs for this sample, number above FACS plot is the mean ± SD, n=6). **(I)** Flow cytometry analysis bone marrow lineages labeled with GFP at week 9 (n=6) and week 13 (n=6). **(J)** Fluorescent images of ileal sections from Abcg2^MCM/+^ R26^GFP/+^ mice 72 hours after a single intraperitoneal tamoxifen injection with higher magnification image to the right**. (K)** Fluorescent images of ileal sections from Abcg2^MCM/+^ R26^GFP/+^ mice 4 weeks after a single intraperitoneal tamoxifen injection with higher magnification image to the right. Scale bars: 50 µm; Statistical significance obtained by student’s t-test; ** *P* < 0.01 and *** *P* < 0.001.

### Bone marrow and intestinal SPCs are enriched for endogenous stem cells that give rise to terminally differentiated cells in vivo

Next, we lineage-traced bone marrow SPCs to determine whether they are enriched for endogenous hematopoietic stem cells(Goodell et al., 1996). Bone marrow SPCs were accurately identified with flow cytometry using negative control samples incubated with Verapamil, a non-specific ABC-transporter inhibitor that blocks the side population phenotype (Figures 1F and 1G). In bone marrow of Abgc2^MCM/+^ R26^GFP/+^ mice injected with tamoxifen, 66% of SPCs (Figure 1H) and 65.7 ± 16.8% (n=5) LSK (Lineage marker^−^, Sca-1^+^, Kit^+^) hematopoietic stem cells were labeled with GFP. Moreover, 92.6 ± 8.5% (n=3) of LSK cells within the side population gate were GFP-labeled. To evaluate the contribution of labeled bone marrow SPCs to hematopoiesis, we assessed GFP-labeling of differentiated bone marrow lineages over a four-week chase period. Again, no GFP-labeling of bone marrow cells was observed in vehicle-injected mice (Figure S2A). Ter119^+^ erythroid cells, which actively express *Abcg2* (29), were extensively labeled at both week 9 and week 13 (Figure 1I and Figure S2). Labeling of all other bone marrow lineages that do not express *Abcg2* (Zhou et al., 2001) increased significantly over the four-week chase period highlighting the contribution of endogenous SPCs to hematopoiesis (Figure 1I, Figures S2B and S2C). The level of labeling observed was much higher than a previous attempt to lineage-trace bone marrow SPCs, where only 2-3% of bone marrow SPCs, LSK cells and differentiated lineages were labeled up to a year after tamoxifen induction(Fatima, Zhou, & Sorrentino, 2012). This previous model inefficiently labeled *Abcg2*-expressing cells most likely because it used an IRES-dependent approach to drive translation of both Abcg2 and CreERT2 from the same mRNA. Our knock-in genetic strategy efficiently labeled *Abcg2*-expressing cells, because we replaced the ATG containing exon of the *Abcg2* gene with MerCreMer cDNA. With this approach, we showed that SPCs enrich for endogenous hematopoietic stem cells that are critical for bone marrow homeostasis.

Similar to isolated bone marrow SPCs, intestinal SPCs enrich for stem cells that give rise to all four intestinal lineages in cell culture(Dekaney et al., 2005; von Furstenberg et al., 2014). To determine whether *Abcg2*-expressing intestinal SPCs enrich for endogenous stem cells required for intestinal homeostasis, we evaluated GFP-labeling in the ileum of Abgc2^MCM/+^ R26^GFP/+^ mice after a single injection of tamoxifen (Figure 1C). Three days after a single tamoxifen injection, there was extensive labeling of enterocytes throughout the villi with minimal labeling of cells in the crypts (Figure 1J). In contrast, four weeks after a single injection of tamoxifen, there was widespread labeling of intestinal cells in a striated pattern that extended from the base of crypts to the tips of villi (Figure 1K). Given the high turnover rate of intestinal epithelium and the pattern of GFP-labeling observed in the ileum(Darwich, Aslam, Ashcroft, & Rostami-Hodjegan, 2014), it was clear that long-term intestinal stem cells were labeled and lineage-traced in vivo. Taken together, these studies show that *Abcg2*-expressing SPCs enrich for endogenous stem cells that are integral for adult bone marrow and intestinal homeostasis.

### Cardiac side population cells are efficiently labeled

After establishing the ability of the side population phenotype to identify endogenous stem cells in organs with continuous cellular turnover, we evaluated the role of SPCs in the heart, one of the least regenerative organs in the body. In Abgc2^MCM/+^ R26^GFP/+^ mice, cSPCs accounted for 1.5% of non-cardiomyocytes (Figures 2A and 2B), which was consistent with previous studies from several independent research groups(Hierlihy et al., 2002; Pfister et al., 2008; Yellamilli & van Berlo, 2016). Identified cSPCs were efficiently labeled with 73% expressing GFP (Figure 2C). To evaluate whether labeled cSPCs give rise to cardiac cells, we measured GFP-labeling of non-cardiomyocytes over a four-week chase period using flow cytometry and immunohistochemical staining. There was extensive labeling of cardiac endothelial cells, which are known to express *Abcg2*(Higashikuni et al., 2010), with a small, but significant increase in labeling over the four-week chase period (Figures 2D and 2E). Due to the high level of baseline endothelial cell labeling, we could not assess whether cSPCs give rise to endothelial cells. Interestingly, only a minor fraction of residential cardiac CD45^+^ cells were GFP-labeled (Figures 2F and 2G) although there was extensive bone marrow labeling (Figure 1I). These results might reflect differences in labeling of embryonic and adult-derived macrophages residing in the heart(Epelman et al., 2014). Other non-cardiomyocyte lineages were infrequently labeled (Figures 2F–2L) with overall less than 1% of non-CD31^+^, non-CD45^+^ cells being labeled with the genetic lineage-tracing approach, which is consistent with the level of cSPC labeling observed (Figures 2A-2C).

**Figure 2.**
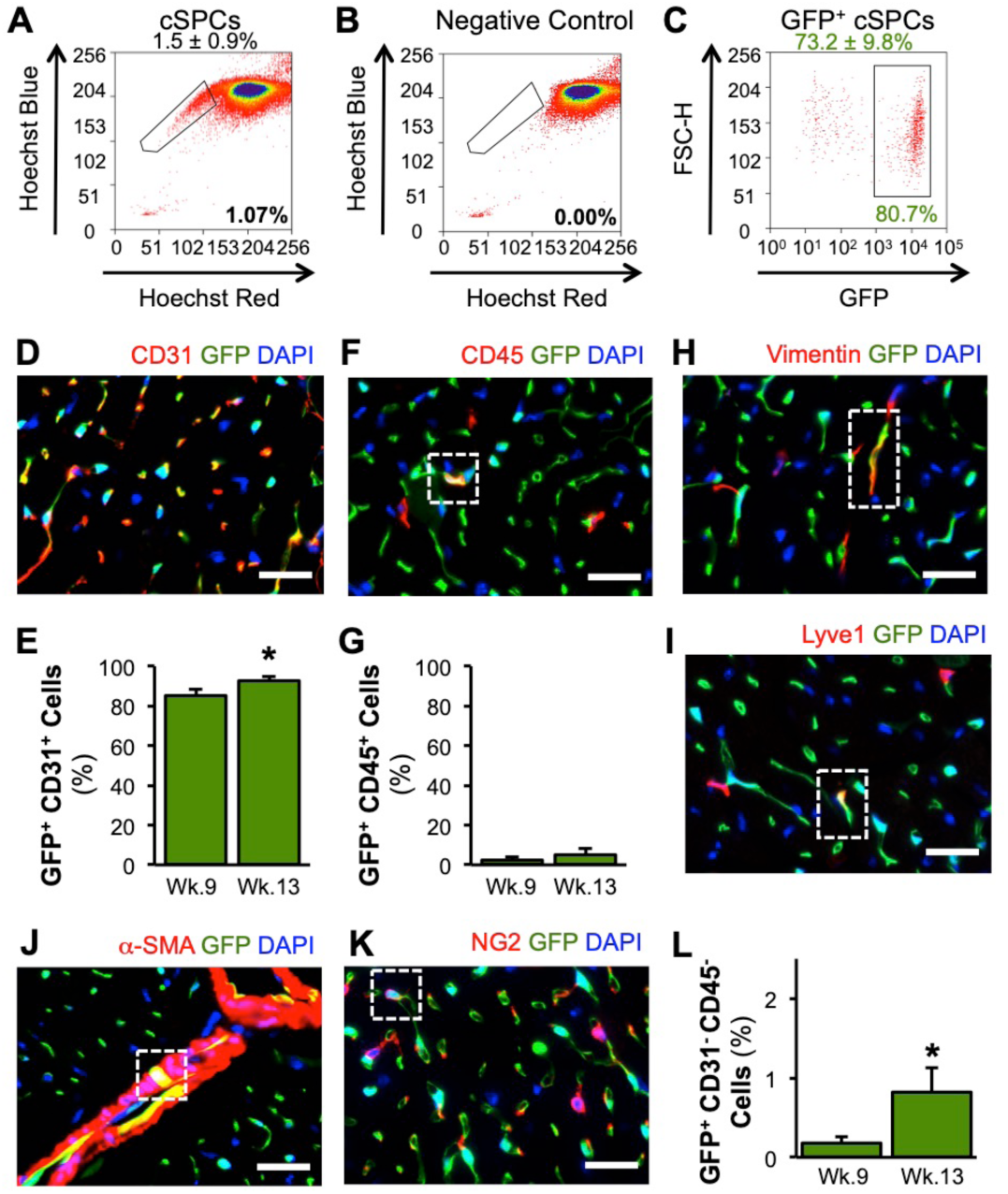
Labeling of cSPCs and non-cardiomyocytes in vivo in the uninjured heart. **(A)** Flow cytometry plot of cSPCs in Abcg2^MCM/+^ R26^GFP/+^ mice treated with tamoxifen (box to the left of the main population, number within FACS plot is the percent of cSPCs for this sample, number above indicates mean ± SD, n=7). **(B)** Corresponding flow cytometry plot of cSPCs stained with Verapamil, which serves as a negative control to ensure accurate identification of SPCs (n=7). **(C)** Flow cytometry plot of GFP fluorescence of cSPCs (number within FACS plot is the percent of GFP^+^ cSPCs for this sample, number above FACS plot is the mean ± SD, n=7). **(D)** Representative immunofluorescence image of CD31 staining on cardiac sections from tamoxifen-injected Abcg2^MCM/+^ R26^GFP/+^ mice. **(E)** Flow cytometry analysis of the percent of CD31^+^ non-cardiomyocytes labeled with GFP at week 9 and week 13 (mean ± SD, n=4). **(F)** Representative immunofluorescence image of CD45 staining on cardiac sections. **(G)** Flow cytometry analysis of the percent of CD45^+^ non-cardiomyocytes labeled with GFP at week 9 and week 13 (mean ± SD, n=4). **(H)** Representative immunofluorescence image of Vimentin staining on cardiac sections. **(I)** Representative immunofluorescence image of Lyve1 staining on cardiac sections. **(J)** Representative immunofluorescence image of α-smooth muscle actin staining on cardiac sections. **(K)** Representative immunofluorescence image of NG2 chondroitin sulfate proteoglycan staining on cardiac sections. **(L)** Flow cytometry analysis of the percent of CD31^−^ CD45^−^ non-cardiomyocytes labeled with GFP at week 9 and week 13 (mean ± SD, n=5). Scale bars: 50 µm; Statistical significance was obtained by student’s t-test; * *P* < 0.05

### Cardiac side population cells contribute to cardiomyocyte labeling during cardiac homeostasis

We observed GFP-labeling of cardiomyocytes in cardiac sections and from adult cardiomyocyte isolations (Figures 3A and 3B). To evaluate to what extent labeled cSPCs contribute to cardiomyocyte labeling, we measured GFP-labeling of isolated adult cardiomyocytes over a four-week chase period and found a five-fold increase from 0.18% to 0.84% (Figures 3C–3F). Although we did not observe active ABCG2-expression in adult cardiomyocytes with immunohistochemistry (Figure 1D), we performed a more rigorous experiment to assess if cardiomyocyte labeling arose from *Abcg2*-expression by cardiomyocytes. We evaluated GFP-labeling of isolated adult cardiomyocytes 72 hours and four weeks after a single tamoxifen injection (Figure 3G). Active expression of *Abcg2* by cardiomyocytes would result in GFP labeling immediately after the tamoxifen injection, while cSPC contribution to cardiomyocytes should give rise to increased labeling over time. At the 72-hour time point, only 0.03% of cardiomyocytes expressed GFP, which increased to 0.74% four weeks later (Figure 3H). These results provided more concrete evidence that lineage-traced cSPCs contribute to cardiomyocyte labeling, and were not the result of cardiomyocyte expression of *Abcg2*.

**Figure 3.**
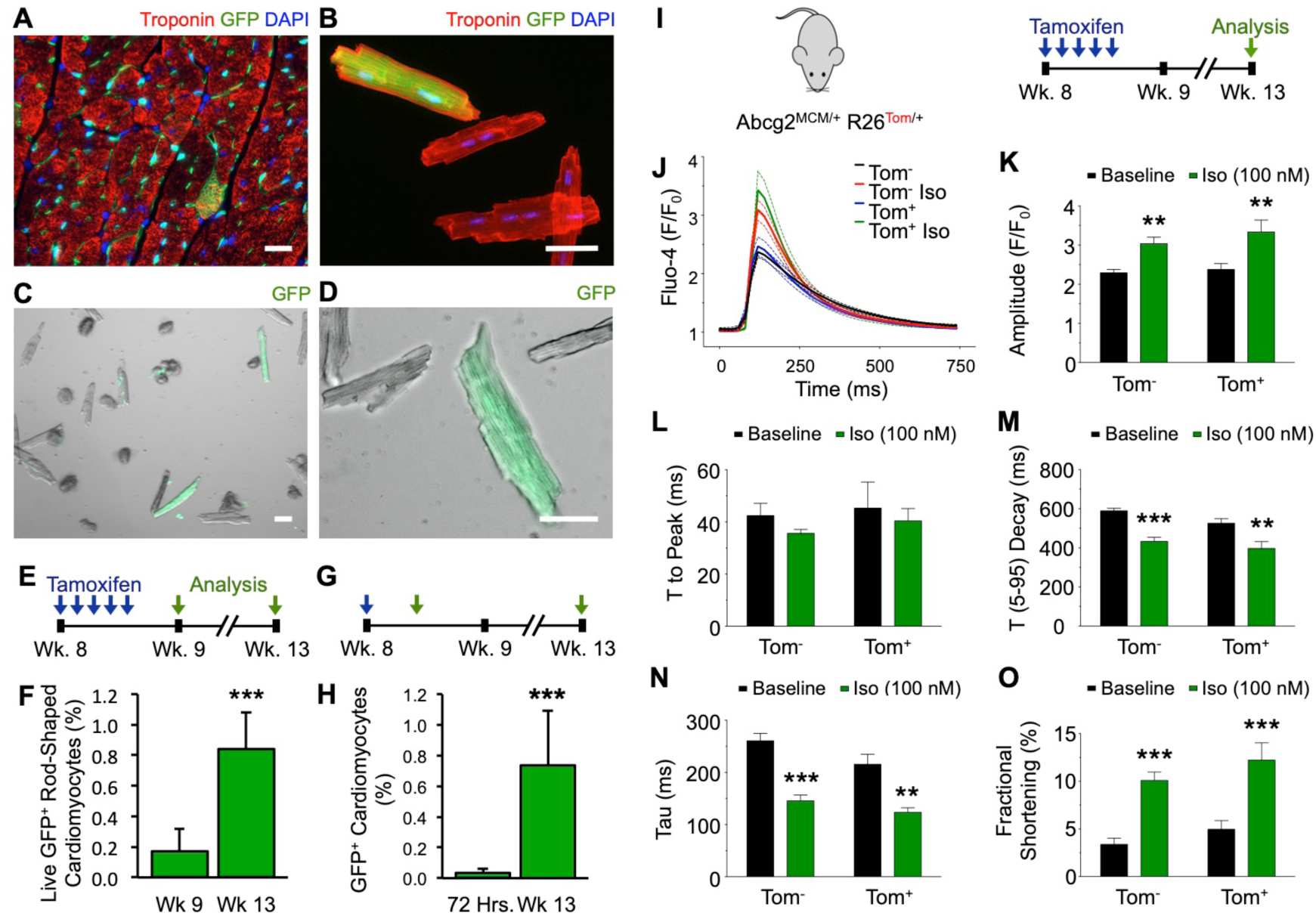
cSPCs contribute to cardiomyocyte labeling in vivo in the uninjured heart. **(A)** Immunofluorescence image of Troponin I staining and native fluorescence of GFP on cardiac sections from tamoxifen-injected Abcg2^MCM/+^ R26^GFP/+^ mice. **(B)** Immunofluorescence image of Troponin T staining and native fluorescence of GFP in fixed, isolated adult cardiomyocytes. **(C)** Image of native GFP fluorescence of live, isolated adult cardiomyocytes used to quantify GFP-labeling in tamoxifen-injected Abcg2^MCM/+^ R26^GFP/+^ mice. **(D)** Higher-magnification image of the native GFP fluorescence of live, isolated adult cardiomyocytes. **(E)** Experimental timeline used to evaluate cardiomyocyte GFP-labeling 72-hours (green arrow) and 4-weeks (green arrow) after 5 consecutive daily tamoxifen injections (blue arrows). **(F)** Quantification of total live, rod-shaped, isolated adult cardiomyocytes expressing GFP at week 9 (n=5) and week 13 (n=7). **(G)** Experimental timeline used to evaluate cardiomyocyte GFP-labeling 72-hours and 4 weeks after single tamoxifen injection. **(H)** Quantification of cardiomyocyte GFP-labeling from cardiac sections 72 hours (n=6) and 4 weeks (n=6) after a single tamoxifen injection. **(I)** Schematic illustrating Abcg2^MCM/+^ R26^Tom/+^ mice and timeline used for calcium dynamics measurements. **(J)** Representative Fluo-4 traces of individual cardiomyocytes. **(K)** Amplitude of pacing-induced cytosolic calcium traces (1Hz) of Tomato^−^ and Tomato^+^ cardiomyocytes with and without Isoproterenol stimulation (Iso 100nM). **(L)** Time (T) to peak Fluo-4 fluorescence of pacing-induced cytosolic calcium traces. **(M)** Time from 5% to 95% of decay of Fluo-4 fluorescence of pacing-induced cytosolic calcium traces. **(N)** Time constant of Ca^2+^ transient decay (Tau) of pacing-induced cytosolic calcium traces. **(O)** Fractional shortening of individual Tomato^−^ and Tomato^+^ cardiomyocytes with and without Isoproterenol stimulation. Scale bars: 50 µm; Statistical significance was obtained by student’s t-test to compare two groups and 2-way ANOVA to compare multiple groups; ** *P* < 0.01, *** *P* < 0.001

To assess the function of labeled cardiomyocytes, we measured cytoplasmic calcium (cCa^2+^) dynamics and contractility in isolated adult cardiomyocytes under baseline conditions and in response to isoproterenol stimulation. We observed normal Ca^2+^ cycling, fractional shortening, and adrenergic responsiveness of labeled cardiomyocytes with no difference in cCa^2+^ dynamics and contractility between labeled and unlabeled cardiomyocytes (Figures 3I– 3O). These data demonstrate that lineage-traced *Abcg2*-expressing cells contribute to labeling of fully-functional adult cardiomyocytes in the uninjured heart.

### Bone marrow cells and endothelial cells do not give rise to cardiomyocyte labeling

Since bone marrow SPCs were extensively labeled in Abcg2^MCM/+^ R26^GFP/+^ mice and have been reported to give rise to cardiomyocytes(Deb et al., 2003; Jackson et al., 2001), we next evaluated the contribution of lineage-traced bone marrow SPCs to labeled cardiomyocytes. We generated bone marrow chimeras by transplanting irradiated Myh7^Cre/+^ R26^tdTomato/+^ mice with bone marrow hematopoietic stem cells (HSCs) isolated from Abcg2^MCM/+^ R26^GFP/+^ mice (Figure 4A). We waited ten weeks after transplantation to inject chimeric mice with tamoxifen to allow sufficient time for transplanted HSCs to engraft and reconstitute the bone marrow niche (Figure 4B). After a four-week chase period, labeling of bone marrow cells in chimeric mice was comparable to bone marrow labeling in Abcg2^MCM/+^ R26^GFP/+^ mice at week 13 (Figures 1H and 1I), confirming successful engraftment and lineage-tracing of bone marrow SPCs (Figure 4C). In the hearts of chimeric mice, we rarely observed GFP-labeled cardiomyocytes (0.02 ± 0.03%, Figure 4D). All GFP-labeled cardiomyocytes also expressed tdTomato, indicating that they arose from fusion of GFP-labeled bone-marrow-derived cells with preexisting tdTomato-labeled cardiomyocytes (Figure 4D). These transplantation studies show that bone marrow cells are not the source of cardiomyocyte labeling in Abcg2^MCM/+^ R26^GFP/+^ mice.

**Figure 4.**
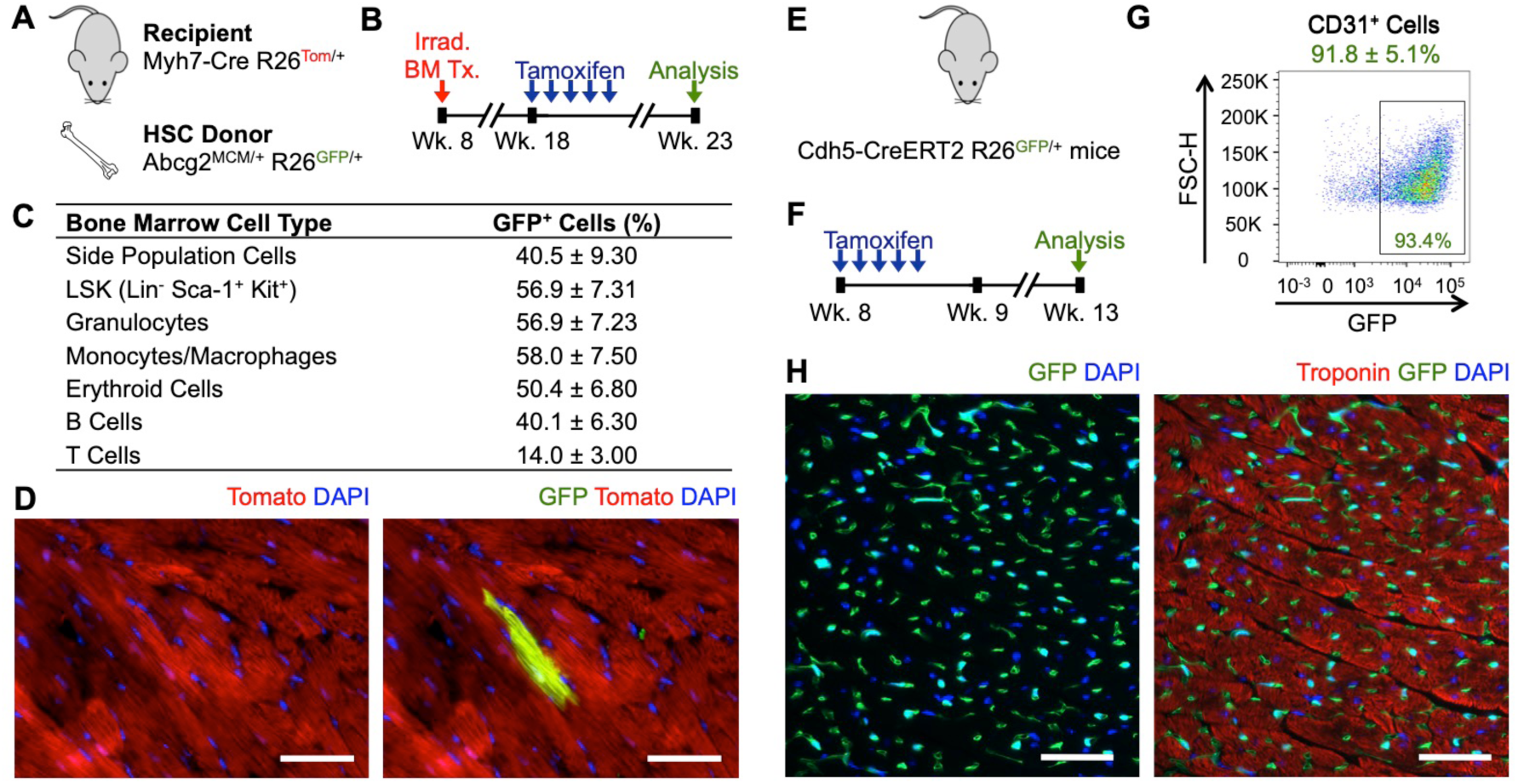
Labeled bone marrow and endothelial cells do not contribute to cardiomyocyte labeling. **(A)** Genetics of hematopoietic stem cell (HSC) donor and recipient mice used to generate bone marrow chimeric Myh7-CreR26^Tom/+^ mice with Abcg2^MCM/+^ R26^GFP/+^ bone marrow. **(B)** Bone marrow transplantation and tamoxifen injection timeline utilized in bone marrow transplantation experiments. **(C)** Assessment of GFP-labeling of bone marrow lineages in chimeric mice four weeks after the fifth tamoxifen injection represented as the percentage of a specific cell type that expresses GFP (mean ± SD, n=5). **(D)** Representative fluorescent image of GFP^+^ and Tom^+^ cardiomyocytes in bone marrow chimeric mice four weeks after the fifth tamoxifen injection. 0.02 ± 0.031% (mean ± SD, n=5) of ventricular cardiomyocytes were labeled with GFP. All GFP-labeled cardiomyocytes also expressed Tomato. **(E)** Genetics of BAC-Cdh5-CreERT2 R26^GFP/+^ mice used to evaluate endothelial cell contribution to cardiomyocyte lineage tracing. **(F)** Experimental timeline used to induce recombination in endothelial cell studies. **(G)** Flow cytometry plot of GFP-labeling of CD31^+^ non-cardiomyocytes in BAC-Cdh5-CreERT2 R26^GFP/+^ mice (number within FACS plot is the percent of GFP^+^ CD31^+^ cells for this sample, number above FACS plot is the mean ± SD n=4). **(H)** Fluorescent image of Troponin I staining and native fluorescence of GFP in BAC-Cdh5^CreERT/+^ R26^GFP/+^ mice at week 13. No GFP^+^ cardiomyocytes were observed in Cdh5^CreERT/+^ R26^GFP/+^ mice (n=5). Scale bars: 50 µm.

Cardiac endothelial cells have also been shown to contribute to cardiomyocytes in the adult heart(Fioret, Heimfeld, Paik, & Hatzopoulos, 2014; Matsuura et al., 2004). To assess whether GFP-labeled endothelial cells contribute to cardiomyocyte-labeling, we evaluated cardiomyocyte labeling in BAC-Cdh5^CreER/+^ mice, an endothelial-specific(Okabe et al., 2014), tamoxifen-inducible Cre transgenic mouse model (Figures 4E and 4F). Four weeks after tamoxifen treatment, endothelial cells were efficiently labeled with GFP in BAC-Cdh5^CreER/+^ R26^GFP/+^ mice (Figure 4G). Despite efficient labeling of endothelial cells, no GFP-labeled cardiomyocytes were identified, indicating that endothelial cells do not account for the observed cardiomyocyte labeling in Abcg2^MCM/+^ R26^GFP/+^ mice (Figure 4H). Our results demonstrate that *Abcg2*-expressing cSPCs, not bone marrow or endothelial cells, contribute to cardiomyocyte labeling under homeostatic conditions.

### Cardiac injury increases cSPC contribution to cardiomyocyte labeling

Next, we assessed how different types of cardiac injury influence the contribution of cSPCs to cardiomyocyte labeling. Since cSPCs and endogenous cardiac regeneration are both activated in response to MI(Hsieh et al., 2007; Senyo et al., 2013; Unno et al., 2012; Yellamilli & van Berlo, 2016), we permanently ligated the left coronary artery in Abcg2^MCM/+^ R26^GFP/+^ mice and analyzed GFP-labeling of cardiomyocytes from histological sections 4 weeks later (Figure 5A-5B). Cardiomyocyte labeling was significantly higher 4 weeks after MI compared to 4 weeks after sham operation (Figures 5C–5E). To ensure that the cardiomyocyte labeling we observed did not arise from cardiomyocytes expressing ABCG2 following MI injury, we stained sections from wild type mice 72 hours after MI injury. We did not detect any cardiomyocytes that expressed ABCG2 within the border zones or remote regions of MI-injured hearts (Figures S3A–S3C). Additionally, we found that cardiac endothelial cells were extensively labeled throughout MI-injured hearts with minimal labeling of fibroblasts and hematopoietic cells (Figures 5F and 5G, and Figure S3D).

**Figure 5.**
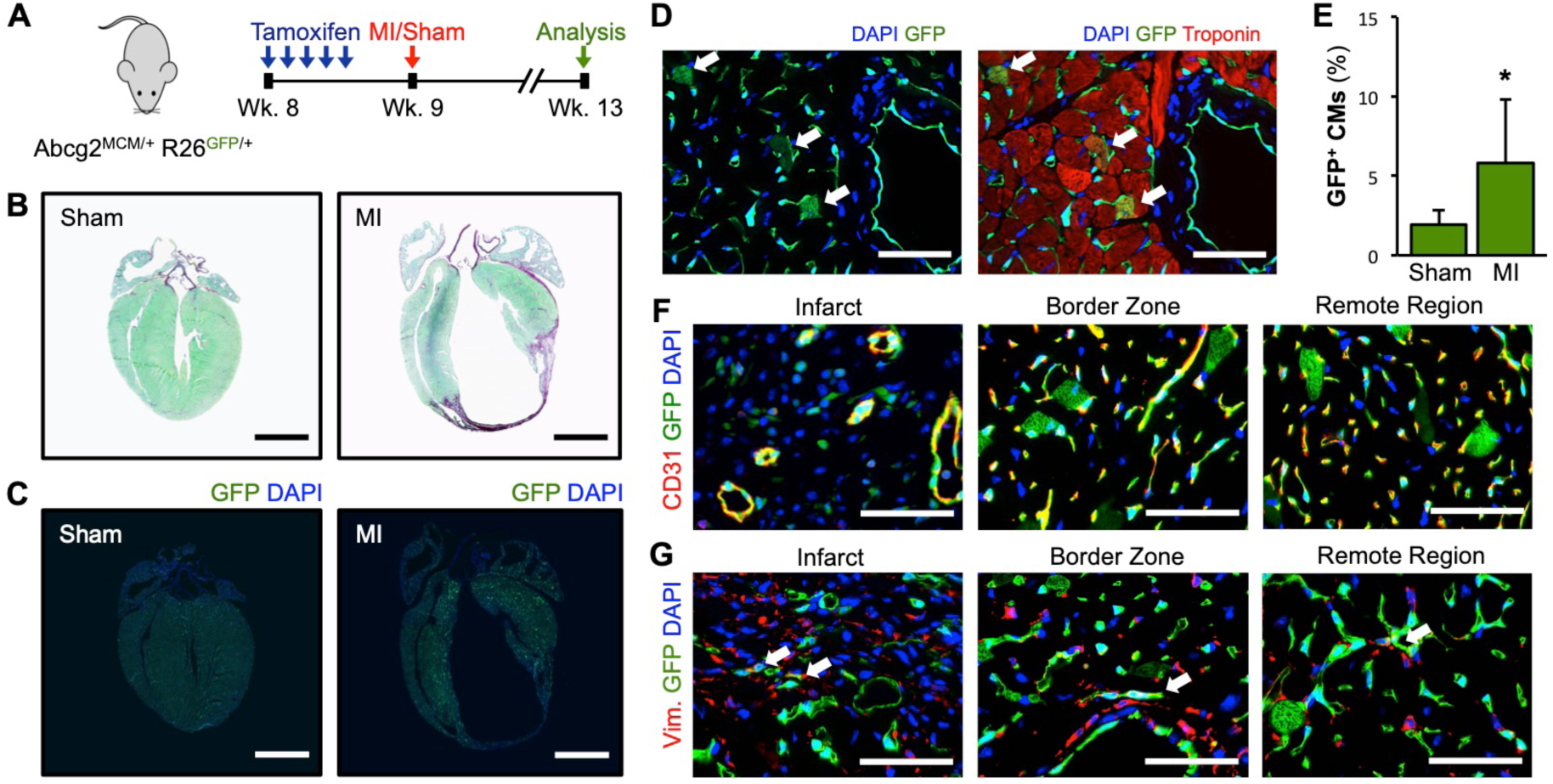
Myocardial ischemic injury increases lineage-traced cardiomyocytes. **(A)** Experimental timeline used to evaluate lineage-tracing of cardiomyocytes after MI in Abcg2^MCM/+^ R26^GFP/+^ mice. 72 hours after the final tamoxifen injections were given, MI or sham operations were performed. Cardiomyocyte labeling was assessed 4 weeks later (green arrow). **(B)** Brightfield images of Sirius Red Fast Green stained cardiac sections from sham and MI-operated Abcg2^MCM/+^ R26^GFP/+^ mice. **(C)** Images of native GFP fluorescence of cardiac sections from sham and MI-operated Abcg2^MCM/+^ R26^GFP/+^ mice. **(D)** Immunofluorescence image of Troponin I staining of cardiac sections from an MI-operated Abcg2^MCM/+^ R26^GFP/+^ mouse. **(E)** Quantification of percent cardiomyocytes expressing GFP (white arrows) on cardiac sections from sham-operated (n=7) and MI-operated (n=8) Abcg2^MCM/+^ R26^GFP/+^ mice (mean ± SD). **(F)** Fluorescent images of CD31 staining to identify endothelial cells in the infarct, border zone and remote regions on histological sections from Abcg2^MCM/+^ R26^GFP/+^ mice four weeks following MI. **(G)** Fluorescent images of Vimentin (Vim.) staining to identify fibroblasts in the infarct, border zone and remote regions on histological sections from Abcg2^MCM/+^ R26^GFP/+^ mice four weeks after MI. White arrows highlight cells that express both GFP and Vimentin. Scale bars: 2 mm (B and C) and 50 µm (D, F and G); Statistical significance was obtained by student’s t-test; * *P* < 0.05.

We also assessed how acute isoproterenol-induced cardiac injury(Brooks & Conrad, 2009; Wallner et al., 2016) impacted cardiomyocyte labeling (Figures S4A–S4C). Similar to the response following MI, there was a higher percentage of cardiomyocytes labeled in isoproterenol-injected mice compared to saline-injected controls (Figure S4A–S4C). These data demonstrate that the contribution of cSPCs to cardiomyocytes is further enhanced in response to different types of cardiac injury.

### Cardiac side population cells fuse with preexisting cardiomyocytes to stimulate cardiomyocyte cell cycle reentry

Finally, we assessed the cellular mechanisms by which *Abcg2-*expressing cSPCs contributed to cardiomyocyte labeling. In addition to directly differentiating into cardiomyocytes, transplanted cardiac progenitor cells have been shown to fuse with preexisting cardiomyocytes(Oh et al., 2003). To assess whether the observed cardiomyocyte labeling arises from direct differentiation of cSPCs or fusion with cSPCs, we cross-bred Abcg2^MCM/+^ mice with R26^mTom-mGFP/mTom-mGFP^ reporter mice (Figure 6A). In Abcg2^MCM/+^ R26^mTom-mGFP/+^ mice, membrane-bound tdTomato (mTom) is expressed in all cells prior to recombination and membrane-bound GFP (mGFP) is expressed after recombination. Seventy-two hours after tamoxifen injections, all mGFP labeled cardiomyocytes also expressed mTom. After a 4-week chase period, 85.3% of GFP-labeled cardiomyocytes remained mTom positive indicating that they likely arose from fusion events, while 14.6% did not express mTom indicating that they potentially arose from direct differentiation of cSPCs to cardiomyocytes (Figures 6B and 6C). Importantly, we did not detect differences in cross-sectional area of mGFP^+^mTom^−^, mGFP^+^mTom^+^, or mGFP^−^mTom^+^ cardiomyocytes (data not shown).

**Figure 6.**
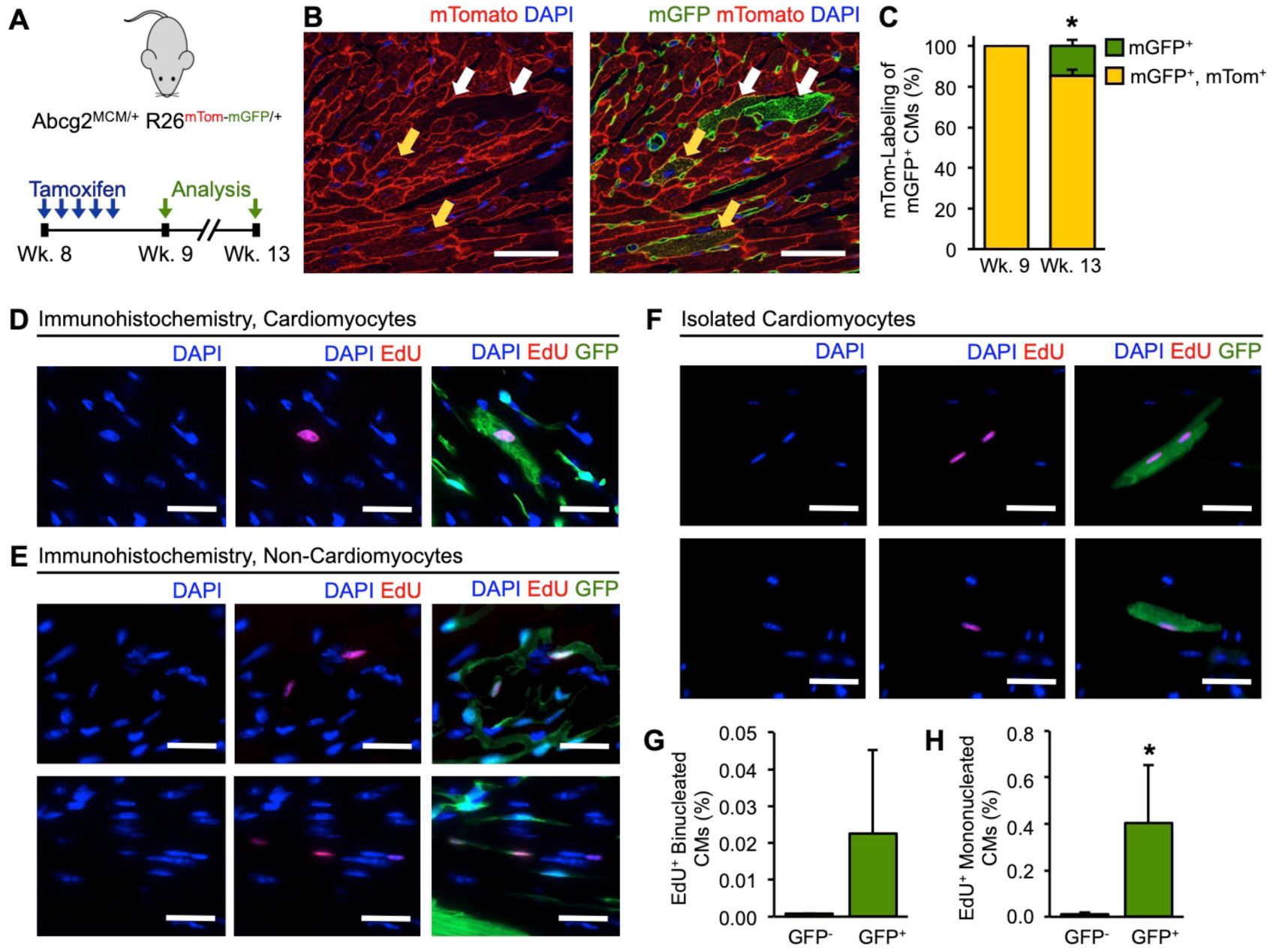
Cardiac SPCs fuse with preexisting cardiomyocytes to stimulate cardiomyocyte cell cycle reentry. **(A)** Experimental design to assess cellular fusion in Abcg2^MCM/+^ R26^mTom-mGFP/+^ mice. **(B)** Fluorescent image of cardiomyocytes labeled with only mGFP (white arrows) or with both mGFP and mTom (yellow arrows). **(C)** Quantification of the percentage of mGFP^+^ cardiomyocytes that are mTom^+^ (yellow) or mTom^−^ (green) at week 9 (n=5) and week 13 (n=4). **(D)** Fluorescent image of an Edu^+^ GFP^+^ cardiomyocyte on histological sections from an Abcg2^MCM/+^ R26^GFP/+^ mouse co-injected with tamoxifen and EdU. **(E)** Fluorescent image of Edu^+^ GFP^+^ non-cardiomyocytes on histological sections from an Abcg2^MCM/+^ R26^GFP/+^ mouse co-injected with tamoxifen and EdU. **(F)** Fluorescent images of binucleated and mononucleated EdU^+^ GFP^+^ isolated adult cardiomyocytes from Abcg2^MCM/+^ R26^GFP/+^ mice. **(F)** Quantification of the percent GFP^−^ mononucleated, rod-shaped cardiomyocytes that is EdU^+^ compared to the percent of GFP^+^ mononucleated, rod-shaped cardiomyocytes that is EdU^+^ at week 13 (mean ± SD, n=4). **(G)** Quantification of the percent GFP^−^ binucleated, rod-shaped cardiomyocytes that is EdU^+^ compared to the percent of GFP^+^ binucleated, rod-shaped cardiomyocytes that is EdU^+^ at week 13 (mean ± SD, n=4). Scale bars: 25 µm; Statistical significance was obtained by student’s t-test; * *P* < 0.05.

To more stringently determine whether a proportion of lineage-traced cardiomyocytes were newly formed, we injected EdU with tamoxifen into Abcg2^MCM/+^ R26^GFP/+^ mice and assessed EdU incorporation four weeks later (Figure S5A). We observed EdU incorporation in nuclei of GFP-labeled cardiomyocytes and non-cardiomyocytes on histological sections (Figures 6D and 6E). To accurately identify and quantify EdU-incorporation and nucleation of GFP-labeled cardiomyocytes, we again injected EdU and tamoxifen into Abcg2^MCM/+^ R26^GFP/+^ mice but quantified isolated adult cardiomyocytes stained for EdU and DAPI (Figures 6F, S5B and S5C). There was no difference in the nucleation of GFP^+^ and GFP^−^ cardiomyocytes (Figures S5D– S5G). We observed EdU incorporation in 0.0012% of all isolated adult cardiomyocytes, which translated to an overall cardiomyocyte DNA synthesis rate of 0.77% per year (Figure S5B and S5C), which is consistent with previous studies(Senyo et al., 2013). Importantly, we observed a significant enrichment of GFP-labeling in EdU^+^ cardiomyocytes with 21% of EdU^+^ cardiomyocytes being GFP-labeled while only 0.7% of all cardiomyocytes were GFP-labeled (Figure 6F and Figure S5C). We also measured nucleation of all EdU^+^ cardiomyocytes and found a trend towards a higher percentage of binucleated EdU^+^ cardiomyocytes labeled with GFP compared to unlabeled binucleated EdU^+^ cardiomyocytes (Figure 6G) and a significantly higher percentage of mononuclear EdU^+^ GFP^+^ cardiomyocytes (Figure 6H). These data clearly show that Abcg2-expressing SPCs contribute to cardiomyocyte cell cycle reentry, and likely to new cardiomyocyte formation.

## DISCUSSION

Evidence supporting the existence of endogenous cardiac regeneration, the heart’s innate ability to form new cells, opened up opportunities for regenerating the failing heart. Previously, the adult mammalian heart was believed to be a post-mitotic organ that could not generate new cardiomyocytes. However, in the past ten years, studies in mice and humans have convincingly demonstrated that a small number of new cardiomyocytes are formed after birth(Bergmann et al., 2009; Bergmann et al., 2015; Hsieh et al., 2007; Senyo et al., 2013). Unfortunately, the number of cardiomyocytes generated is not sufficient to compensate for those lost during the progression of heart failure. Future therapies that enhance endogenous cardiac regeneration could improve clinical outcomes when used in conjunction with current heart failure therapies.

Many different pre-clinical approaches to enhance cardiac regeneration are actively being pursued(Eschenhagen et al., 2017; Garbern & Lee, 2013; Laflamme & Murry, 2011). In recent years, there has been increased focus on accomplishing this by stimulating cardiomyocyte proliferation(Leach et al., 2017; Nakada et al., 2017). Moreover, recent evidence suggests that presumed cardiac progenitor cells, which were thought to have the advantage of giving rise to non-cardiomyocytes and cardiomyocytes, both of which are integral to normal cardiac function(Bollini et al., 2014; Garbern & Lee, 2013), are generally incapable of generating cardiomyocytes(Li et al., 2018; van Berlo & Molkentin, 2014, 2016). Growing evidence has accumulated that *Kit*-expressing cells, the most extensively studied group of proposed cardiac progenitor cells, have a negligible contribution to endogenous cardiac regeneration(He et al., 2017; Sultana et al., 2015; van Berlo et al., 2014). For these reasons, it is important that we evaluate the potential role of cells that are distinct from cardiac *Kit*-expressing cells. One such population of cells is cSPCs, which do not express *Kit* at the mRNA or protein level(Mouquet et al., 2005; Noseda et al., 2015). Furthermore, high-throughput transcriptional analysis comparing cardiac *Kit*-expressing cells to cSPCs demonstrated the striking difference in expression profiles between these two populations of cells(Dey et al., 2013).

Here, we used a genetic lineage-tracing approach to investigate whether the side population phenotype can be used to identify endogenous cardiac non-cardiomyocytes that give rise to cardiomyocytes during cardiac homeostasis and in response to cardiac injury. We used *Abcg2*-expression to lineage-trace SPCs since studies have shown that ABCG2 is essential for the side population phenotype but is not required for the function of SPCs as stem cells under homeostatic conditions(Martin et al., 2004; Zhou et al., 2002; Zhou et al., 2001). In bone marrow SPCs, ABCG2 is the only ABC transporter responsible for the side population phenotype(Zhou et al., 2002). In the adult heart, ABCG2 and p-glycoprotein, which is encoded by *Mdr1a* and *Mdr1b*, are both responsible for the side population phenotype(Pfister et al., 2008). When *Abcg2* is knocked out, mice are completely viable and have no defects in bone marrow or cardiac homeostasis(Pfister et al., 2008; Zhou et al., 2002). Therefore, we used an *Abcg2*-driven lineage-tracing mouse model to trace SPCs in vivo to determine whether they are enriched for endogenous stem cells in the heart and other tissues throughout the body, even though Abcg2 is not exclusive to side population cells. Currently, no unique marker for side population cells is known, and future studies will need to evaluate a more precise marker.

With this mouse model, we found that cSPCs contribute to cardiomyocyte labeling during cardiac homeostasis with an increase in this contribution following both isoproterenol-induced and MI injury. We showed that lineage-traced bone marrow cells and endothelial cells do not contribute to cardiomyocyte labeling, which fits with previous literature(Alvarez-Dolado et al., 2003; Mouquet et al., 2005; Murry et al., 2004; Nygren et al., 2004). More importantly, we found that cardiac SPCs fuse with preexisting cardiomyocytes and induce cell cycle reentry of lineage-traced cardiomyocytes. Our findings build on previous reports of cardiomyocyte fusion with non-cardiomyocytes. In cell culture, cardiomyocytes have the potential to fuse with endothelial cells and fibroblasts(Matsuura et al., 2004). These findings were corroborated by genetic lineage tracing of cardiomyocytes, which showed that almost 2% of cardiomyocytes in the adult heart had undergone fusion with an unidentified non-cardiomyocyte population of cells(Hsieh et al., 2007). Subsequent studies identified multiple cell types, including circulating bone marrow cells, Sca-1+ cells, and c-kit+ cells that have the potential to fuse with cardiomyocytes(Alvarez-Dolado et al., 2003; He et al., 2017; Nygren et al., 2004; Oh et al., 2003; van Berlo et al., 2014; Wu et al., 2015). Our results extend these findings by providing evidence that residential Abcg2-expressing non-cardiomyocytes fuse with pre-existing cardiomyocytes at rates, which account for most cardiomyocyte fusion events reported by Hsieh et al.(Hsieh et al., 2007).

While fusion and proliferation are thought of as distinct processes, there is growing evidence that cardiomyocyte fusion can trigger cell cycle entry and induce cardiomyocyte proliferation. When cardiomyocytes are co-cultured with endothelial cells and fibroblasts, they fuse with surrounding non-cardiomyocytes, re-enter the cell cycle, and begin to proliferate(Matsuura et al., 2004). In a recent study, it was shown that transient membrane fusion is essential for cardiomyocyte proliferation in adult zebrafish(Sawamiphak, Kontarakis, Filosa, Reischauer, & Stainier, 2017). More recently, a study in mice showed that non-cardiomyocytes expressing *Twist2* have the ability to fuse with preexisting cardiomyocytes; however, the relationship between fusion and cardiomyocyte proliferation was not investigated(Min et al., 2018). Our study provides additional evidence that residential non-cardiomyocytes fuse with cardiomyocytes. More importantly, we provide evidence with our lineage-tracing mouse model, which shows that proliferative cardiomyocytes can arise from fusion of *Abcg2*-expressing cSPCs with cardiomyocytes. When we consider our findings along with recently published studies, they suggest that fusion between cardiomyocytes and non-cardiomyocytes regulates endogenous cardiomyocyte proliferation in the adult mammalian heart.

An alternative interpretation of the data we have presented here is that a small number of proliferative cardiomyocytes could express *Abcg2*. While we did not observe ABCG2-expression by cardiomyocytes with immunohistochemistry, we cannot conclusively rule out that *Abcg2* is expressed by a small number of cardiomyocytes. In two studies, researchers found that diseased human hearts expressed ABCG2 at the protein level; however, the appropriate negative controls were not available for these human samples to check the specificity of the methodologies used(Alfakir et al., 2012; Meissner et al., 2006; Solbach et al., 2008). Additionally, these studies did not determine whether cardiomyocytes or non-cardiomyocytes expressed ABCG2 in their samples(Alfakir et al., 2012; Meissner et al., 2006; Solbach et al., 2008). In our studies, we used cardiac tissue from *Abcg2* knockout mice to rigorously evaluate the specificity of four commercially available ABCG2 antibodies and found that only one specifically bound ABCG2. Despite the care we took to accurately assess ABCG2 expression in cardiomyocytes, it is impossible to prove that cardiomyocytes never express *Abcg2*. Our study with single tamoxifen injections added a more rigorous approach to carefully assess whether our genetic lineage-tracing model is activated within cardiomyocytes. Since we observed minimal levels of recombination in cardiomyocytes 72-hours after a single injection of tamoxifen, we concluded that the significant increase in cardiomyocyte labeling during the four-week chase period arose from cSPCs. Similarly, after 5 tamoxifen injections, the increase in cardiomyocyte labeling was likely due to fusion between cSPCs and cardiomyocytes.

In this study, we used the side population phenotype to identify a population of residential non-cardiomyocytes that have the ability to fuse with preexisting cardiomyocytes and trigger cardiomyocyte cell cycle reentry. Our data suggests that this mechanism might account for ~20% of proliferative cardiomyocytes in the adult heart during cardiac homeostasis. Further research is needed to characterize fusogenic cells identified by the side population phenotype and to understand their relationship with other non-cardiomyocytes(Min et al., 2018). Moving forward, careful work will be needed to understand how fusion triggers cell-cycle reentry in cardiomyocytes and whether this mechanism can be targeted for future regenerative therapies.

## MATERIALS AND METHODS

### Experimental Mouse Models

All animal procedures were performed conform the NIH guidelines and approved by the University of Minnesota Institutional Animal Care and Use Committee. No human subjects or human material was used. Gene targeting of the murine Abcg2 locus to insert a complementary DNA encoding Cre recombinase flanked by a mutated estrogen receptor was done by standard gene targeting. A targeting vector containing ampicillin resistance and a diphtheria toxin A cassette was used to insert homology arms upstream and downstream of the ATG start codon containing second exon of Abcg2, the cDNA for MerCreMer cloned in-frame with the ATG start codon of Abcg2, as well as a frt site flanked neomycin selection cassette through recombineering. A linearized targeting vector was electroporated into embryonic stem (ES) cells and correctly targeted clones were identified by PCR and Southern blot analysis. ES cell aggregation with 8-cell embryos was used to generate chimeric mice. Germline transmitting chimeras were cross-bred with Rosa26-Flpe mice (B6.129S4-Gt(ROSA)26Sortm1(FLP1)Dym/RainJ) to delete the neomycin selection cassette. We generated experimental animals by cross-breeding Abcg2^MCM/+^ mice to previously modified FVB.Cg-Gt(ROSA)26Sor^tm1(CAG-lacZ,EGFP)Glh^/J (van Berlo et al., 2014), B6.129(Cg)-Gt(ROSA)26Sor^tm4(ACTB-tdTomato,-EGFP)Luo^/J or B6.Cg-*Gt(ROSA)26Sor*^*tm14(CAG-td<u>Tomato</u>)Hze*^/J mice (all reporter mice were purchased from the Jackson Laboratory). A previously published BAC transgenic Cdh5 CreERT2 mouse line was cross-bred to R26^GFP/GFP^ mice(Okabe et al., 2014). A previously published Myosin Heavy Chain 7Cre mouse line was cross-bred to B6.Cg-*Gt(ROSA)26Sor*^*tm14(CAG-td<u>Tomato</u>)Hze*^/J mice(Parsons et al., 2004). PCR genotyping of Abcg2^MCM/+^ was performed with the following primers: wild type fw: 5’-tcaaagtgctggtatctgtgttga-3’, wild type rev: 5’-catgaattgaagtatccacagcaa-3’, mut fw: 5’-ggtgggacatttgagttgct-3’, mut rev: 5’-catatgtacaacaacatgaattgaagtatcc-3’, MerCreMer fw: 5’-ggcgttttctgagcatacct-3’, MerCreMer rev: 5’-ctacaccagagacggaaatcc-3’. Previously published Abcg2 knockout mice were used to verify antibody specificity(Zhou et al., 2002). Both males and female mice were used in all experiments.

### Chemicals

To induce Cre-mediated recombination, 8-week-old mice were injected intraperitoneally with 2 mg of tamoxifen for 5 consecutive days. DNA synthesis was measured by labeling with 5-Ethynyl-2’-deoxyuridine (EdU). The timing and frequency of intraperitoneal injections is shown in the experimental schematics, where every arrow indicates one intraperitoneal injection of 2.5 mg of EdU. For isoproterenol studies, mice were given subcutaneous 100 mg/kg injections below the loose skin overlying their scapulae once a day, for five consecutive days starting 72 hours after the final tamoxifen injection was given. Vehicle control mice were subcutaneously injected with a similar volume 0.9% sodium chloride solution.

### Myocardial infarction surgery

Seventy-two hours after the final tamoxifen injection, mice were anesthetized with 3% isoflurane, intubated via intratracheal intubation and maintained on 2.5% isoflurane throughout the surgery. A parasternal thoracotomy was performed, followed by permanent ligation of the left coronary artery just below the left atrial auricle using 7-0 silk suture(Gundewar, Calvert, Elrod, & Lefer, 2007). After confirming ligation by visual observation of myocardial blanching distal to the suture; the musculature and skin were sequentially closed in layers and mice were allowed to recover on a heating pad. For sham-operated mice, the same steps were completed except for ligation of the left coronary artery. Mice were administered subcutaneous buprenorphine SR-LAB prior to the start of surgery. Aseptic technique was used for all surgical procedures.

### Isolation and FACS analysis of bone marrow side population cells and lineages

Bone marrow cells were isolated and analyzed for side population phenotype and presence of lineage markers using published protocols(Ergen, Jeong, Lin, Challen, & Goodell, 2013). Bone marrow was isolated from bilateral femora and tibiae. For side population analysis, bone marrow cells were stained with 5 µg/mL Hoechst 33342 for 90 minutes in a 37°C water bath. As a negative control, bone marrow cells were similarly stained with 5 µg/mL Hoechst 33342 but 50 µM verapamil was added for 90 minutes in a 37°C water bath. Hoechst 33342-stained cells were subsequently stained with a biotinylated lineage antibody panel followed by staining with streptavidin conjugated to Alexa Fluor™ 647, α-Sca-1 PE antibody and α-Kit APC antibody for Lineage^−^Sca-1^+^Kit^+^ (LSK) analysis. For bone marrow lineages, freshly isolated bone marrow cells were stained with either α-CD3ε biotin, α-CD11b biotin, α-CD45R biotin, α-Ly6G and Ly6C biotin or α-Ter119 biotin followed by streptavidin conjugated to Alexa Fluor 647. Before FACS analysis, propidium iodide was added to all samples at a final concentration of 2 µg/mL for live cell/dead cell discrimination. Bone marrow side population cell data were acquired using the MoFlo XDP flow cytometer cell sorter and analyzed using the accompanying Summit™ software.

### Bone marrow transplantation

Transgenic Myosin Heavy Chain 7 ^Cre/+^ mice cross-bred with R26^Tomato/+^ mice were conditioned with 11.0 Gy total body irradiation one day before transplantation. Bone marrow was isolated from Abcg2^MCM/+^ R26^GFP/+^ mice that had not received tamoxifen injections. Hematopoietic stem cells were enriched using the EasySep Mouse Hematopoietic Progenitor Cell Enrichment kit (Stemcell Technologies) and 1*10^6^ HSCs were administered via tail vein injection to recipient mice. Ten weeks after transplantation, bone marrow chimeric mice were administered intraperitoneal injections of 2 mg tamoxifen for 5 consecutive days. Four weeks after the final tamoxifen injection, GFP-labeling of bone marrow side population cells and differentiated lineages, and of cardiomyocytes from histological sections were analyzed.

### Isolation and FACS analysis of cardiac side population cells and non-cardiomyocyte lineages

Non-cardiomyocytes were isolated from the heart and analyzed for the side population phenotype and lineage analysis using a previously published protocol(Pfister, Oikonomopoulos, Sereti, & Liao, 2010). Hearts were flushed with ice-cold PBS to remove red blood cells, minced into a fine slurry and digested in a solution containing 2.4 U/mL Dispase II, 0.1% Collagenase B and 2.5 mM calcium chloride for 30 minutes in a 37°C water bath. Next, the digestion solution was triturated, strained through a 70 µm cell strainer followed by a 40 µm cell strainer, and centrifuged at 600 g for 5 minutes at 4°C to pellet non-cardiomyocytes out of the digestion solution. For cardiac side population analysis, non-cardiomyocytes were stained with 1.5 µg/mL Hoechst 33342 for 90 minutes in a 37°C water bath. As a negative control, non-cardiomyocytes were also stained with 1.5 µg/mL Hoechst 33342 and 50 µM verapamil for 90 minutes in a 37°C water bath. For non-cardiomyocyte lineages, freshly isolated non-cardiomyocytes were stained with either α-CD45 PE or α-CD31 APC. Before FACS analysis, propidium iodide was added to all samples at a final concentration of 2 µg/mL for live cell/dead cell discrimination. Cardiac side population cell data were acquired using a MoFlo™ XDP flow cytometer cell sorter and analyzed using the accompanying Summit software. Non-cardiomyocyte lineage data were acquired using a BD FACSAria II flow cytometer cell sorter and analyzed using FlowJo v10 software application.

### Tissue processing for histology

All tissues for histology were fixed with 4% paraformaldehyde for 3 hours at room temperature with gentle rocking, washed with PBS and incubated in a 30% sucrose solution (w/v in PBS) overnight at 4°C. After tissues sunk down to the bottom of specimen tubes, they were trimmed, embedded in O.C.T. and frozen over a slurry of dry ice and Isopentane. Hearts were cut and embedded in a four-chamber orientation. Blocks were cut into 5 µm sections and were mounted onto Fisherbrand Superfrost Plus Microscope slides. Blocks and slides were stored at −80°C until further use.

### Histological stains, image acquisition and analysis

To identify non-cardiomyocytes and cardiomyocytes on histological sections, cryosections were stained with primary antibodies. Briefly, slides were air-dried for 5 minutes, washed with 0.1% Triton X-100 (v/v in PBS), blocked for 1 hour at room temperature in antibody-specific blocking solution, and incubated with primary antibodies overnight at 4°C. The following day, slides were washed, incubated with secondary antibodies and DAPI and washed before coverslips were mounted on the slides using Vectashield mounting medium. Antibodies used include Abcg2 (BXP-53 Santa-Cruz, rat monoclonal 1:400), CD31 (Lifespan Biosciences LS-B4737, rabbit polyclonal 1:20), Vimentin (Abcam ab45939, rabbit polyclonal 1:50), CD45 (R&D Systems, goat polyclonal 1:20), α-SMA (Abcam ab5694, rabbit polyclonal 1:150), Lyve1 (Abcam ab14917, rabbit polyclonal 1:250), NG2 (Millipore AB5320, rabbit polyclonal 1:150), Troponin I (Abcam ab47003, rabbit polyclonal 1:100). For EdU staining, the Click-iT Plus EdU Alexa Fluor Imaging Kit was used and nuclei were counter stained with DAPI. All immunohistochemically stained slides were imaged using a Zeiss Axio Imager M1 Upright Microscope.

To quantify GFP-labeling of cardiomyocytes on histological sections, cryosections were stained with Wheat Germ Agglutinin conjugated to Texas Red-X and DAPI. Ten separate images were taken with a 20X objective throughout the left ventricle using the Zeiss Axio Imager M1 Upright Microscope. The number of GFP-labeled cardiomyocytes was counted for each field-of-view and quantification of GFP-labeled cardiomyocytes was calculated as the percent of GFP^+^ over total cardiomyocytes.

For Sirius Red/Fast Green staining, cryosection slides were fixed with pre-warmed Bouin’s fixative at 55°C for 1 hour, stained with 0.4% Fast Green (v/v in picric acid) for 10 minutes at room temperature followed by 0.1% Sirius Red (v/v in picric acid) for 30 minutes at room temperature. After each stain, slides were washed with running tap water to remove excess dye. Coverslips were mounted with Permount Mounting Medium. Slides were then imaged using a Huron Technologies TISSUEscope LE brightfield slide scanner available at the University Imaging Center at the University of Minnesota. Slide images were viewed using the free HuronViewer software. Frozen sections from Abcg2^MCM/+^ R26^mTmG/+^ were stained with DAPI and imaged using a Nikon C2 and Nikon A1 confocal microscope at the University Imaging Centers, University of Minnesota.

### Isolation and analysis of adult cardiomyocyte

Adult cardiomyocytes were isolated using a previously published protocol(O’Connell, Rodrigo, & Simpson, 2007). After excision, hearts were cannulated on a gravity-dependent Langendorff perfusion setup and perfused with 2.4 mg/mL Collagenase type 2 solution for 9-11 minutes. Digested hearts were cut into 10-12 pieces and gentle triturated to release individual cardiomyocytes. Cardiomyocytes were pelleted by centrifuging digestion solution at 19 g for 5 minutes. The total numbers of rod-shaped and rounded-up cardiomyocytes were counted using a Fuchs-Rosenthal counting chamber. The number of GFP-labeled rod-shaped cardiomyocytes was counted by an individual blinded to the experimental conditions using a Zeiss Axio Observer Z1 Inverted Microscope. Quantification of GFP-labeled cardiomyocytes was calculated as the percent of total rod-shaped cardiomyocytes labeled with GFP.

For Troponin T and EdU staining, isolated adult cardiomyocytes were immediately fixed with 4% paraformaldehyde (v/v in PBS) for 5 minutes followed by washing with PBS before EdU and Troponin T staining. For Troponin T staining, cardiomyocytes were washed with 0.1% Triton X-100 (v/v in PBS), blocked for an hour at room temperature with 5% normal goat serum (v/v in PBS), and incubated with Troponin T antibody (Thermo Fisher MS295, mouse monoclonal 1:100) overnight at 4°C. The following day, cardiomyocytes were washed and incubated with Donkey α-mouse antibody Alexa Fluor 568 and DAPI (50 ng/mL). EdU-stained cardiomyocytes were also stained for GFP (Abcam ab290, rabbit polyclonal 1:100) using the same protocol outlined for Troponin T staining. All data from isolated adult cardiomyocytes was acquired using a Zeiss Axio Observer Z1 Inverted Microscope.

### Adult Cardiomyocyte cytosolic calcium (_c_Ca^2+^) transient recordings

Isolated adult cardiomyocytes were loaded with 1 µM Fluo-4 AM and placed in a 37°C chamber on an inverted microscope stage (Zeiss Axio Observer Z1, Sutter DG4 plus: excitation, and Sutter Lambda 10-2: emission and images are acquired using a Photometrics Evolve Delta EMCCD camera) as previously detailed(Luongo et al., 2017). Adult cardiomyocytes were perfused with a physiological Tyrode’s buffer (150 mM NaCl, 5.4 mM KCl, 1.2 mM MgCl_2_, 10 mM glucose, 2 mM sodium pyruvate, and 5 mM HEPES, pH 7.4) containing 2 mM CaCl_2_. Cells were paced at 1 Hz and _c_Ca^2+^ transients continuously recorded and analyzed offline with pCLAMP 10 software. After two minutes of baseline recording, 100 nM isoproterenol was applied by changing the perfusion solution. For _c_Ca^2+^ fluorescence measurements, background fluorescence was subtracted and changes in [Ca^2+^] are expressed as changes of normalized fluorescence, F/F0, where F denotes the maximal fluorescence (peak) and F0 (or F unstimulated), is the resting fluorescence at the beginning of each recording (average fluorescence of the cell 50 ms prior to stimulation). For fractional shortening measurements, isolated adult cardiomyocytes were placed in a heated chamber (37°C) on the stage of an inverted microscope and perfused with a normal physiological Tyrode’s buffer 2 mM CaCl_2_ and paced by field-stimulation at 1 Hz (100 ms pulse duration). Fractional shortening measurements were taken from both ends of each cell using a video edge detection system (Crescent Electronics) and after two minutes of baseline recording, 100 nM isoproterenol (Sigma-Aldrich) was applied by changing the perfusion solution. All data is paired, i.e. all baseline %FS values have accompanying values post-iso treatment.

### Statistical Analysis

Results are reported as mean ± standard deviation. For adult cardiomyocyte cytosolic calcium transient recordings and fractional shortening measurements data are reported as mean ± standard error of the mean. Student’s t-tests were performed to compare two groups and 2-way ANOVA to compare multiple groups. p-values less than 0.05 were considered statistically significant.

## AUTHOR CONTRIBUTIONS

A.Y. and J.H.v.B. contributed to study design, data analysis, and writing the manuscript. A.Y. was involved with all assays and data collection. Y.R. acquired and analyzed MoFlo flow cytometry data. R.T.M. and J.T. generated bone marrow chimeric mice. J.P.L., P.G., S.M., S.R.H., and J.W.E. performed Ca^2+^ transient and fractional shortening experiments. D.J.G. assisted with data interpretation and edited the manuscript.

## ACKNOWLEDGMENTS

The authors would like to thank Xiaodan Wang, Natsumi Nemoto, Chetana Guthikonda, Jessica Shaklee, Wuqiang Zhu, Ingrid Bender and John Calvert for their assistance. This work was supported by the National Institutes of Health (HL112852, HL130072 to J.H.v.B. and MSTP T32GM008244 to A.Y.). J.H.v.B. is supported by The Hartwell Foundation and Minnesota Regenerative Medicine Initiative.

## Supplemental Information

**Figure S1.**
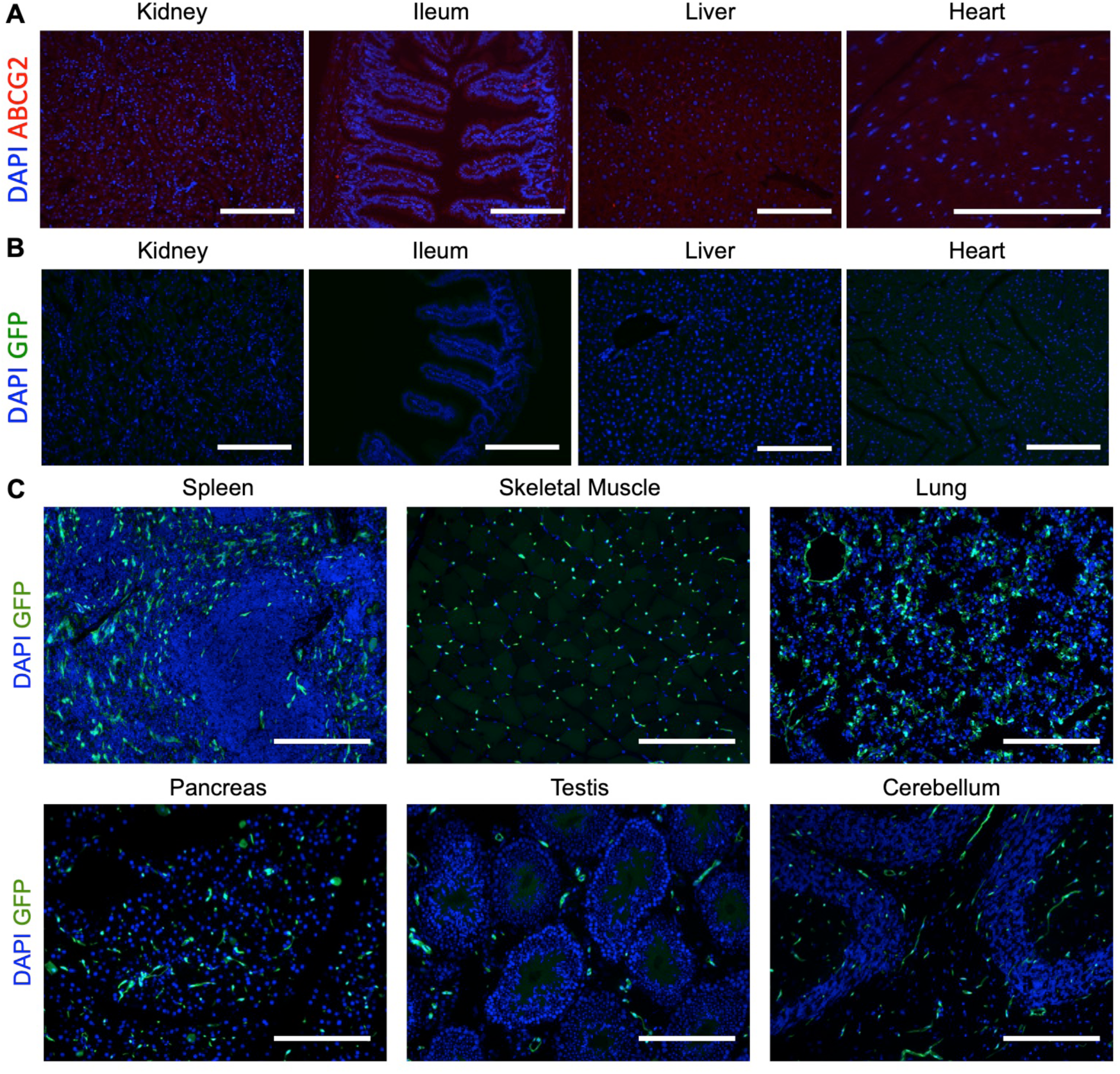
Controls used to validate Abcg2-driven lineage tracing mouse model (Related to Figure 1). **(A)** Fluorescent images of renal, ileal, hepatic and cardiac sections from Abcg2^−/−^ mice stained for ABCG2. **(B)** Fluorescent images of renal, ileal, hepatic and cardiac sections from Abcg2^MCM/+^ R26^GFP/+^ mice injected with corn oil vehicle. **(C)** Images of native GFP fluorescence in indicated organs from tamoxifen-injected Abcg2^MCM/+^ R26^GFP/+^ mice at week 9. Scale bars: 200 µm.

**Figure S2.**
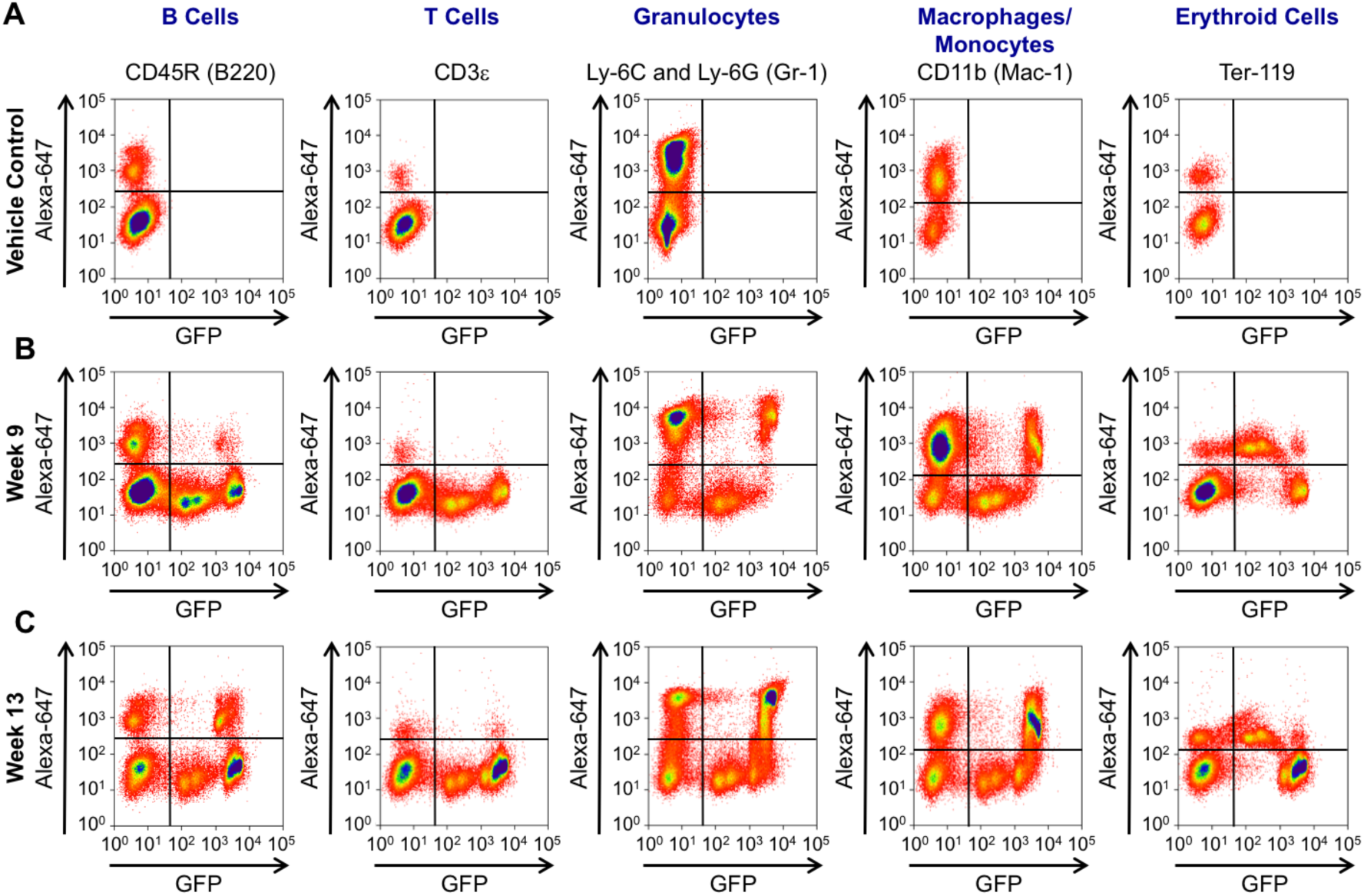
Flow cytometric analysis of GFP-labeled bone marrow lineages isolated from Abcg2^MCM/+^ R26^GFP/+^ mice (Related to Figure 1). **(A)** Flow cytometry plots of GFP fluorescence of differentiated bone marrow lineages from Abcg2^MCM/+^ R26^GFP/+^ mice injected with corn-oil vehicle. **(B)** Flow cytometry plots of GFP fluorescence of differentiated bone marrow lineages from tamoxifen-injected Abcg2^MCM/+^ R26^GFP/+^ mice at week 9. **(C)** Flow cytometry plots of GFP fluorescence of differentiated bone marrow lineages from tamoxifen-injected Abcg2^MCM/+^ R26^GFP/+^ mice at week 13. Quantification of GFP-labeled bone marrow lineages is calculated as the percentage of cells in the upper right quadrant (GFP^+^) over the total number of cells positive for the lineage marker in both the upper right and upper left quadrants (Lineage marker^+^).

**Figure S3.**
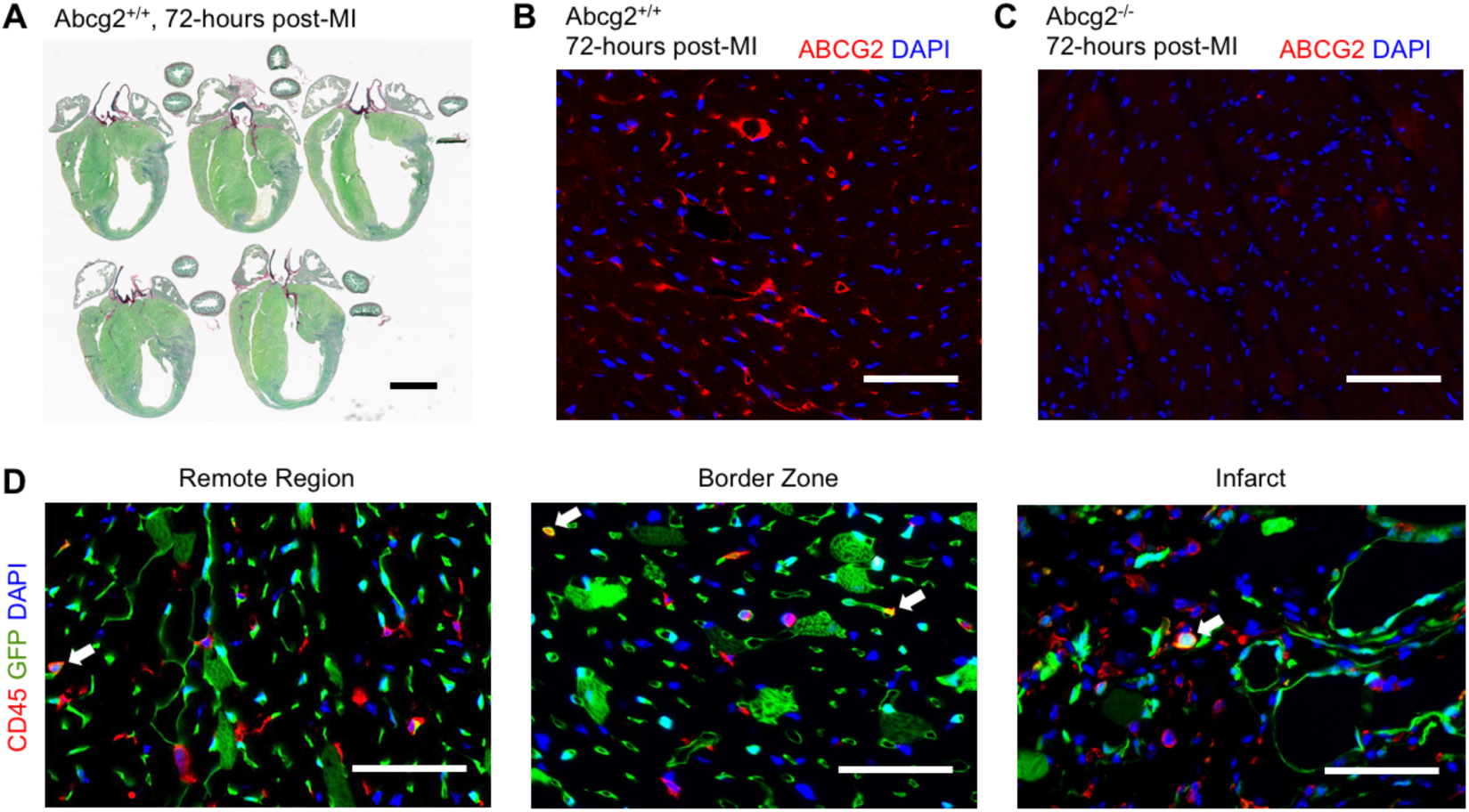
Characterization of ABCG2 expression and non-cardiomyocyte lineage-tracing following myocardial ischemic injury (Related to Figure 5). **(A)** Brightfield images of Sirius Red Fast Green stained cardiac sections 72 hours following MI in wild type (Abcg2^+/+^) mice demonstrating transmural scar formation. **(B)** Fluorescent images of sections harvested from Abcg2^+/+^ mice 72 hours following MI stained for ABCG2 (red) and DAPI (blue). **(C)** Fluorescent images of sections harvested from Abcg2^−/−^ mice 72 hours following MI stained for ABCG2 (red) and DAPI (blue) as control to validate staining in panel **(B). (D)** Fluorescent images of CD45 staining of histological sections from Abcg2^MCM/+^ R26^GFP/+^ mice 4 weeks following MI. White arrows highlight cells that express both GFP and CD45. Scale bars: 50 µm.

**Figure S4.**
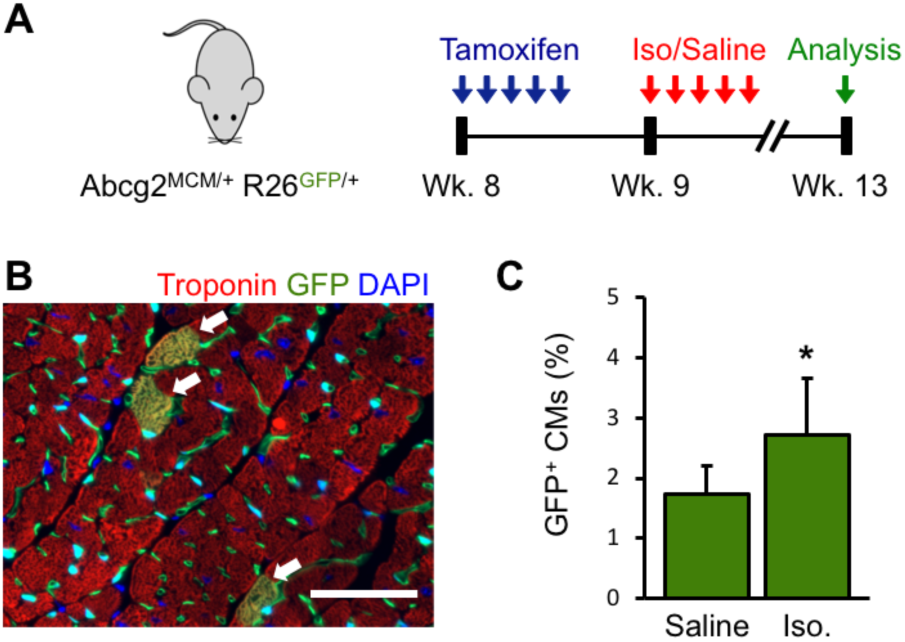
Acute isoproterenol injury increases lineage-traced cardiomyocytes (Related to Figure 5). **(A)** Experimental timeline used to evaluate lineage-traced cardiomyocytes following acute isoproterenol injury in Abcg2^MCM/+^ R26^GFP/+^ mice. 72 hours after the final tamoxifen injection, mice were injected daily with isoproterenol or saline for 5 days. Cardiomyocyte labeling was assessed at week 13, i.e. 4 weeks after isoproterenol or saline injections were given. **(B)** Immunofluorescence image of Troponin I staining (red) on cardiac sections from an isoproterenol-injected Abcg2^MCM/+^ R26^GFP/+^ mouse. **(C)** Quantification of GFP-labeled cardiomyocytes from cardiac sections of saline-injected and isoproterenol-injected Abcg2^MCM/+^ R26GFP/+ mice (mean ± SD, n=8 and 10 resp.). Scale bar: 50 µm; Statistical significance was assessed by Student’s t-test; * P < 0.05

**Figure S5.**
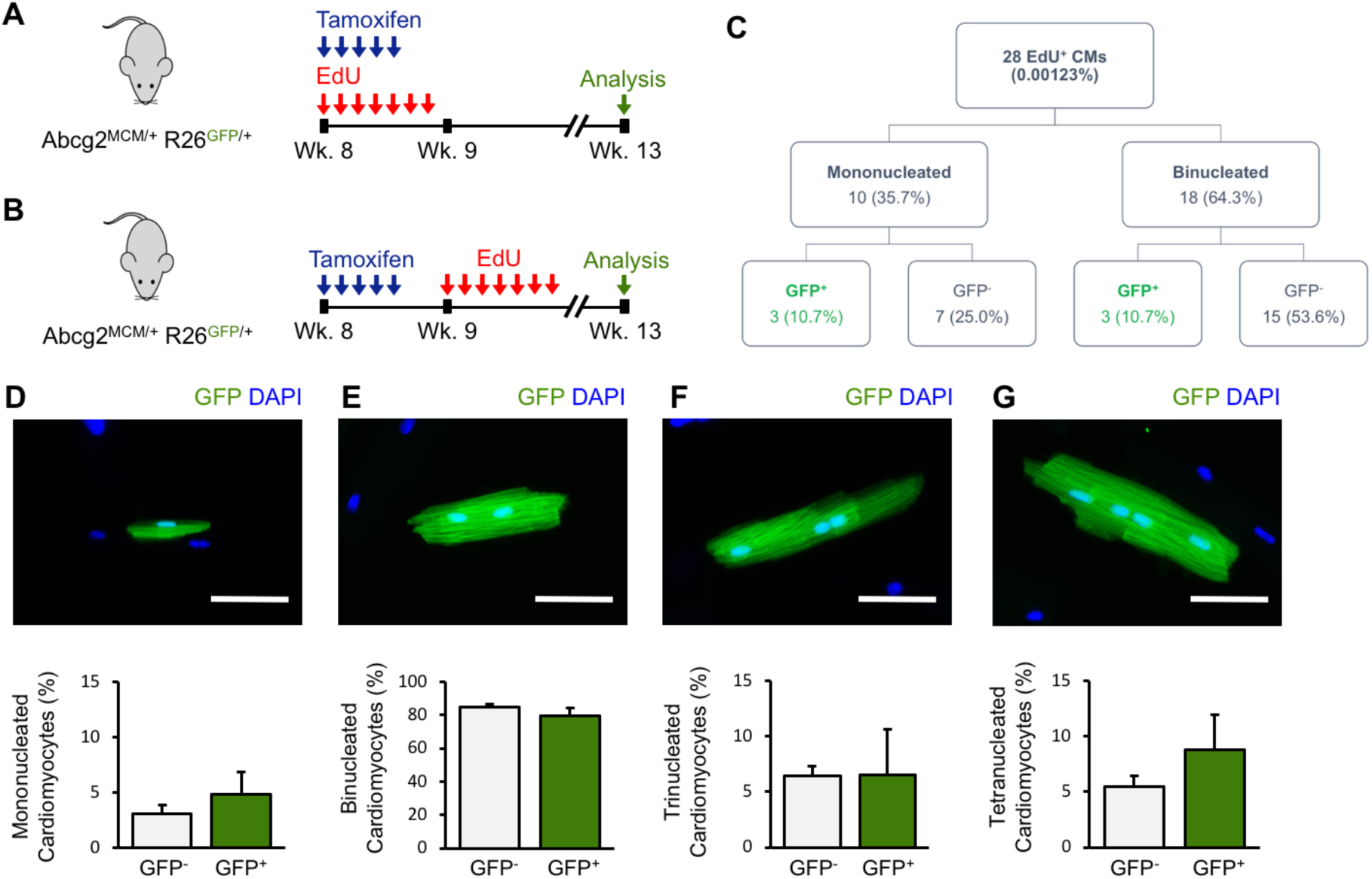
EdU labeling of cardiomyocytes and non-cardiomyocytes from Abcg2^MCM/+^ R26^GFP/+^ mice at week 13 (Related to Figure 6). **(A)** Experimental design to assess proliferation in Abcg2^MCM/+^ R26^GFP/+^ mice co-injected with tamoxifen and EdU, related to Figures 6D and 6E. **(B)** Experimental design to assess cardiomyocyte proliferation in Abcg2^MCM/+^ R26^GFP/+^ mice sequentially injected with tamoxifen then EdU, related to Figures 6F–6H. **(C)** Tree diagram of nucleation and GFP-labeling of EdU^+^ cardiomyocytes (CMs) from Abcg2^MCM/+^ R26^GFP/+^ mice. **(D)** Fluorescent image of a mononucleated GFP^+^ cardiomyocyte and quantification of the percent of GFP^+^ and GFP^−^ cardiomyocytes that are mononucleated. **(E)** Fluorescent image of a binucleated GFP^+^ cardiomyocyte and quantification of the percent of GFP^+^ and GFP^−^ cardiomyocytes that are binucleated. **(F)** Fluorescent image of a trinucleated GFP^+^ cardiomyocyte and quantification of the percent of GFP^+^ and GFP^−^ cardiomyocytes that are trinucleated. **(G)** Fluorescent image of a tetranucleated GFP^+^ cardiomyocyte and quantification of the percent of GFP^+^ and GFP^−^ cardiomyocytes that are tetranucleated. Scale bars: 50 µm; Statistical significance was assessed by Student’s t-test; (mean ± SD; n=4).

### SUPPLEMENTAL EXPERIMENTAL PROCEDURES

#### Myocardial infarction surgery

Seventy-two hours after the final tamoxifen injection, mice were anesthetized with 3% isoflurane, intubated via intratracheal intubation and maintained on 2.5% isoflurane throughout the surgery. A parasternal thoracotomy was performed, followed by permanent ligation of the left coronary artery just below the left atrial auricle using 7-0 silk suture(Gundewar et al., 2007). After confirming ligation by visual observation of myocardial blanching distal to the suture; the musculature and skin were sequentially closed in layers and mice were allowed to recover on a heating pad. For sham-operated mice, the same steps were completed except for ligation of the left coronary artery. Mice were administered subcutaneous buprenorphine SR-LAB prior to the start of surgery. Aseptic technique was used for all surgical procedures.

#### Isolation and FACS analysis of bone marrow side population cells and lineages

Bone marrow cells were isolated and analyzed for side population phenotype and presence of lineage markers using published protocols(Ergen et al., 2013). Bone marrow was isolated from bilateral femora and tibiae. For side population analysis, bone marrow cells were stained with 5 µg/mL Hoechst 33342 for 90 minutes in a 37°C water bath. As a negative control, bone marrow cells were similarly stained with 5 µg/mL Hoechst 33342 but 50 µM verapamil was added for 90 minutes in a 37°C water bath. Hoechst 33342-stained cells were subsequently stained with a biotinylated lineage antibody panel followed by staining with streptavidin conjugated to Alexa Fluor™ 647, α-Sca-1 PE antibody and α-Kit APC antibody for Lineage^−^Sca-1^+^Kit^+^ (LSK) analysis. For bone marrow lineages, freshly isolated bone marrow cells were stained with either α-CD3ε biotin, α-CD11b biotin, α-CD45R biotin, α-Ly6G and Ly6C biotin or α-Ter119 biotin followed by streptavidin conjugated to Alexa Fluor 647. Before FACS analysis, propidium iodide was added to all samples at a final concentration of 2 µg/mL for live cell/dead cell discrimination. Bone marrow side population cell data were acquired using the MoFlo XDP flow cytometer cell sorter and analyzed using the accompanying Summit™ software.

#### Isolation and FACS analysis of cardiac side population cells and non-cardiomyocyte lineages

Non-cardiomyocytes were isolated from the heart and analyzed for the side population phenotype and lineage analysis using a previously published protocol(Pfister et al., 2010). Hearts were flushed with ice-cold PBS to remove red blood cells, minced into a fine slurry and digested in a solution containing 2.4 U/mL Dispase II, 0.1% Collagenase B and 2.5 mM calcium chloride for 30 minutes in a 37°C water bath. Next, the digestion solution was triturated, strained through a 70 µm cell strainer followed by a 40 µm cell strainer, and centrifuged at 600 g for 5 minutes at 4°C to pellet non-cardiomyocytes out of the digestion solution. For cardiac side population analysis, non-cardiomyocytes were stained with 1.5 µg/mL Hoechst 33342 for 90 minutes in a 37°C water bath. As a negative control, non-cardiomyocytes were also stained with 1.5 µg/mL Hoechst 33342 and 50 µM verapamil for 90 minutes in a 37°C water bath. For non-cardiomyocyte lineages, freshly isolated non-cardiomyocytes were stained with either α-CD45 PE or α-CD31 APC. Before FACS analysis, propidium iodide was added to all samples at a final concentration of 2 µg/mL for live cell/dead cell discrimination. Cardiac side population cell data were acquired using a MoFlo™ XDP flow cytometer cell sorter and analyzed using the accompanying Summit software. Non-cardiomyocyte lineage data were acquired using a BD FACSAria II flow cytometer cell sorter and analyzed using FlowJo v10 software application.

#### Isolation and analysis of adult cardiomyocyte

Adult cardiomyocytes were isolated using a previously published protocol(O’Connell et al., 2007). After excision, hearts were cannulated on a gravity-dependent Langendorff perfusion setup and perfused with 2.4 mg/mL Collagenase type 2 solution for 9-11 minutes. Digested hearts were cut into 10-12 pieces and gentle triturated to release individual cardiomyocytes. Cardiomyocytes were pelleted by centrifuging digestion solution at 19 g for 5 minutes. The total numbers of rod-shaped and rounded-up cardiomyocytes were counted using a Fuchs-Rosenthal counting chamber. The number of GFP-labeled rod-shaped cardiomyocytes was counted by an individual blinded to the experimental conditions using a Zeiss Axio Observer Z1 Inverted Microscope. Quantification of GFP-labeled cardiomyocytes was calculated as the percent of total rod-shaped cardiomyocytes labeled with GFP.

